# Chloroplast movements in siphonous macroalgae in response to high light and grazing

**DOI:** 10.64898/2026.05.14.725087

**Authors:** Heta Mattila, Pedro Lopes, Vesa Havurinne, Johannes W. Goessling, Paulo Cartaxana, Sónia Cruz

**Affiliations:** CESAM–Centre for Environmental and Marine Studies, Department of Biology, University of Aveiro, Portugal; Molecular Plant Biology, Department of Life Technologies, University of Turku, Finland

**Keywords:** Chloroplast relocation, Grazing, Herbivory, Kleptoplastic sea slug, Photodamage, Photo-relocation, Seaweed, Ulvophyceae

## Abstract

Fast cytoplasmic streaming enables extensive chloroplast movements in the giant cells of unicellular, siphonous macroalgae. Here, we studied chloroplast movements in two such algae: the Dasycladalean *Acetabularia acetabulum* and the Bryopsidales *Bryopsis* sp.. We hypothesised that chloroplast movements function as a protective avoidance mechanism under excess light, particularly in *Bryopsis* sp., which lacks capacity for fast induction of photoprotective non-photochemical quenching (NPQ) and state transitions. In addition, we also investigated whether chloroplast movements are involved in responses to wounding and herbivory. The movements were studied by light microscopy, photography and pulse modulated chlorophyll *a* fluorescence quenching analysis. Chemical inhibitors of actin polymerization and microtubules assembly were used to confirm that the observed effects were active responses controlled by the cytoskeleton. *A. acetabulum* responded to high light by reversible chloroplast aggregation, probed by macro-imaging; and chemical inhibition of chloroplast movements led to an enhancement of Photosystem II photoinhibition, as probed by the fluorescence parameter F_V_/F_M_. No chloroplast movements were observed in *Bryopsis* sp. in response to high light. In *A. acetabulum*, wounding caused either by cutting or due to feeding by the sap-sucking sea slug *Elysia timida* triggered aggregation of chloroplasts within minutes of incurring the damage. Interestingly, the aggregation also occurred in intact cells away from the cutting site. Furthermore, the addition of media collected from the vicinity of cut algae was sufficient to induce chloroplast aggregation in intact algae, suggesting that water-borne cues or signals triggered the aggregation response in *A. acetabulum*. *Bryopsis* sp., however, responded to cutting by only local chloroplast aggregation. The relevance of chloroplast movements in protection against both abiotic and biotic stressors in *A. acetabulum,* and the potential reasons behind the different defence strategies of the algae, are discussed.

## Introduction

Certain algae, like green macroalgae in the Dasycladales and Bryopsidales clades, can grow to considerable sizes (up to several tens of centimetres) and show complex thallus morphologies, despite being unicellular. In these giant-celled algae, fast cytoplasmic streaming enables substantial chloroplast movements (Schmid & Koop 1983; Menzel & Schliwa 1986; Sugi & Chaen 2003; Ochiai et al. 2024). In the Bryopsidales macroalga *Halimeda macroloba*, for example, a mass movement of chloroplasts occur during growth of a new thallus segment (Larkum et al. 2011). In another Bryopsidales alga, *Caulerpa brachypus*, “macroscopic waves” are observed when chloroplasts move towards the tip of the thallus during the day and away during the night (Saco et al. 2021; Afik et al. 2023). In plants and in most of the studied algae, chloroplast movements are controlled by light receptors (usually by phototropins, phytochromes and/or neochromes) and mediated by actin filaments and myosin motor proteins (Nagai & Fukui 1981; Sugi & Chaen 2003; Wada & Kong 2018; Labuz et al. 2022; Ohtaka & Sekimoto 2023). However, in the Bryopsidales alga *Bryopsis* sp., chloroplast movements appear to depend on microtubules and kinesin motor proteins (Ochiai et al. 2024; Ogawa et al. 2026).

Light, the energy source of photosynthetic organisms, also damages their photosynthetic machinery, especially Photosystem II (PSII), both directly and via the production of harmful reactive oxygen species (Tyystjärvi 2013; Vass 2012; Zavafer 2021; Mattila et al. 2023). In plants and some algae, chloroplasts typically accumulate under low light at the periclinal cell wall, on the top of the cell to maximize light absorption; while under high light, chloroplasts accumulate at the anticlinal (side) cell walls as a light avoidance response (for reviews, see Wada & Kong 2018; Labuz et al. 2022; Ohtaka & Sekimoto 2023). In plants, the avoidance response has been shown to protect chloroplasts from excess light (e.g., Kasahara et al. 2002; Majumdar & Kar 2020).

Phaeophyceae (i.e., brown macroalgae), belonging to the Stramenopiles clade, possess “plant-like” chloroplast movements; *Dictyota dichotoma*, for example, shows periclinal accumulation of chloroplasts under low light (at afternoon or in deep water) and anticlinal accumulation under high light (at noon in shallow water) (Nultsch et al. 1984; Hanelt & Nultsch 1990). Analogously, under low light, the large and flat chloroplasts of the green Zygnematophycean algae align themselves perpendicular to light, while a side-on arrangement is observed under high light (for a review, see Ohtaka & Sekimoto 2023). The siphonous, unicellular yellow-green macroalga (Xanthophyceae; Stramenopiles) *Vaucheria sessilis* shows light-induced chloroplast aggregation (Blatt 1983). In addition to brown and yellow-green macroalgae, chloroplasts in Stramenopile microalgae also respond to high light, as chloroplast aggregation and/or contraction of a single chloroplast (karyostrophy) have been observed in diatoms (e.g., Kiefer 1973; Silkin et al. 2021; Bastos et al. 2025). Chloroplast movements in green macroalgae (Chlorophyta) commonly follow a diurnal rhythm in several species of the clade, like the Ulvophyceaen algae *Acetabularia acetabulum* and *Ulva lactuca* (Dawes & Barilotti 1969; Britz and Briggs 1976; Nultsch et al. 1984; Schmid & Koop 1983; Drew & Abel 1990). Also, the above-described chloroplast movements in the Bryopsidales alga *C. brachypus* continue in the absence of a day/night cycle, suggesting that they are under the control of the circadian clock (Saco et al. 2021; Afik et al. 2023). However, whether these movements in green macroalgae serve a photoprotective function has not been thoroughly explored.

Bryopsidales algae appear to lack the fast, proton-gradient regulated (qE) component of non-photochemical quenching (NPQ) (Christa et al. 2017; Handrich et al. 2017; Iha et al. 2021; Xu et al. 2025). This photoprotective mechanism diminishes production of reactive oxygen species and photoinhibition of both photosystems (PSII and PSI), especially during sudden changes in light intensity (Peers et al. 2009; Allorent et al. 2013; Cantrell & Peers 2017; Roach 2020; Steen et al. 2022). In addition, we recently observed that many Bryopsidales also appear to lack state transitions (Havurinne et al. 2025), a mechanism that balances light excitation between the photosystems. Here, we hypothesized that chloroplast movements function as a compensatory photoprotective mechanism in these algae, particularly as genes related to microtubule-based movements in *Bryopsis corticulans* have been shown to be rapidly upregulated in response to green light (Xu et al. 2025). First, we tested if chloroplast movements protect against PSII photoinhibition by measuring chlorophyll *a* fluorescence in the absence and presence of chloroplast movement inhibitors in two siphonous macroalgae; *Bryopsis* sp. (deficient in qE) and *A. acetabulum* (capable of qE; e.g., Mattila et al. 2025) (see Fig. 1). However, in response to high light, chloroplast movements (in particular, chloroplast aggregation) were observed exclusively in *A. acetabulum*, where they seemed to reduce PSII photoinhibition. Next, pronounced chloroplast aggregation was observed in response to cellular damage caused by cutting or feeding by a “sap-sucking” sea slug herbivore, but again, only in *A. acetabulum*. The relevance of the observed chloroplast movements in response to abiotic and biotic stressors, as well as possible reasons for the different defence strategies between the studied algae, are discussed.

**Figure 1.**
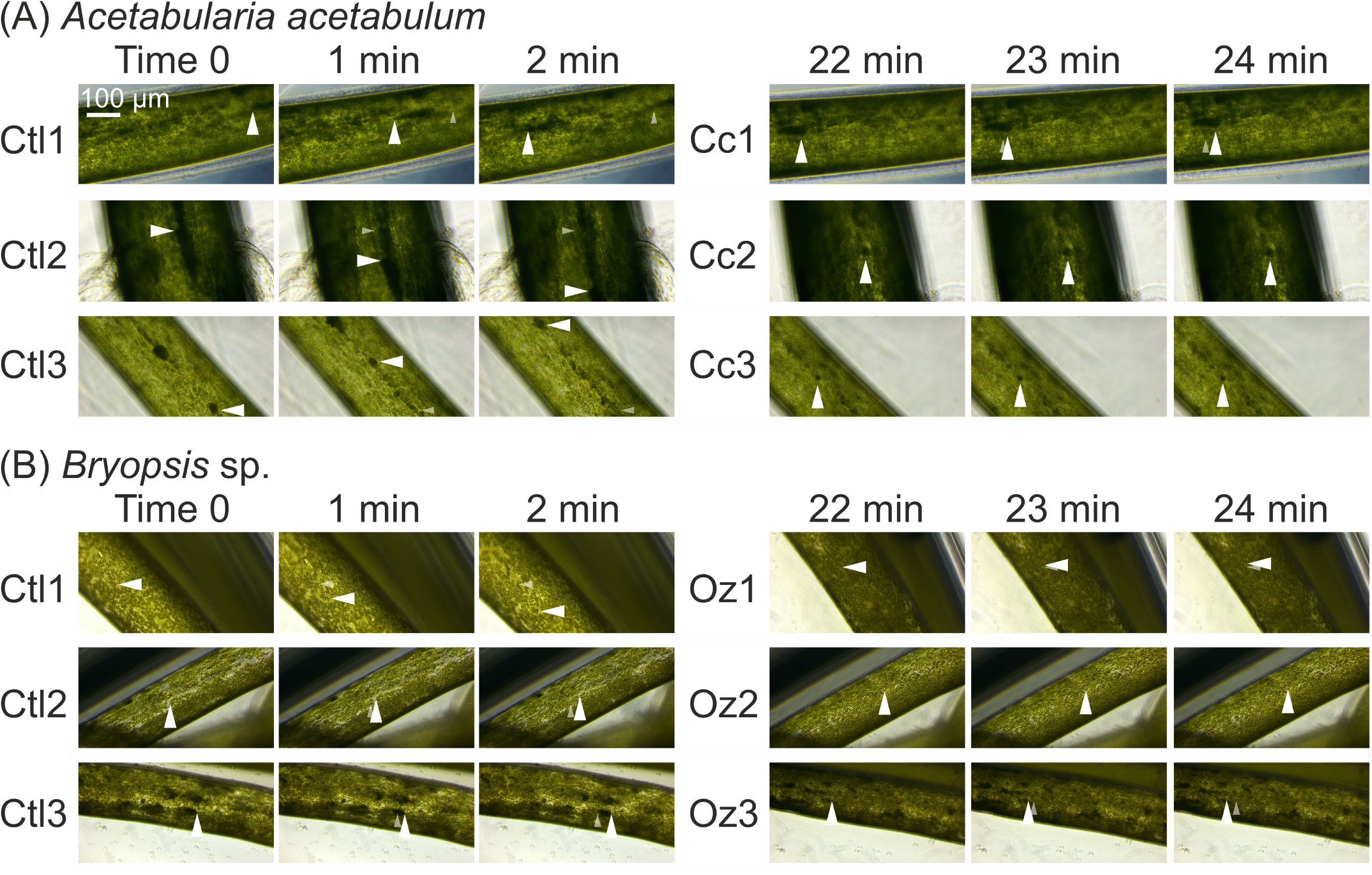
Inhibition of actin (by cytochalasin B) and microtubules (by oryzalin) based cytoplasmic streaming resulted in the loss of chloroplast motility in *Acetabularia acetabulum* and *Bryopsis* sp., respectively. A series of light microscopy images from sections of *A. acetabulum* (A) and *Bryopsis* sp. (B) cells. The first sets of 3 images were captured with intervals of 1 min (Ctl), after which cytochalasin B (Cc; A) or oryzalin (Oz; B) was added. After 20 min of incubation, the imaging was repeated, as indicated. The light of the microscope was kept on during the whole treatment. Three biological replicates (labelled 1–3) are shown. The large, white arrowheads highlight the estimated positions of the same chloroplasts within the 3-image sets. The smaller, transparent arrowheads highlight the initial positions of the chloroplasts (note that in some of the images, the positions of the arrowheads overlap). All images were taken with a 20x objective, and the scale bar is shown in the first image. Brightness and contrast were adjusted equally in each series of 6 images.

## Materials and methods

### Strains and culture conditions

The macroalgae *A. acetabulum* (strain DI1; originally isolated by Diedrik Menzel) and *Bryopsis* sp. (previously annotated as *B. plumosa*; KU-0990; Kobe University Macroalgal Culture Collection, Japan) were cultured in the laboratory at 20–22 °C, under the photosynthetic photon flux density (PPFD) of 40–80 µmol m^−2^ s^−1^ (12 h light/12 h dark), in modified f/2 medium without silica (Andersen 2005) prepared in 35 ppt artificial seawater (Red Sea salt, Red Sea Europe, France). For details, see Havurinne & Tyystjärvi (2020), Cartaxana et al. (2023) and Mattila et al. (2025).

The sea slug *Elysia timida* (originally collected from the Mediterranean Sea, at the coast of Elba, Italy) was cultured in the laboratory at 20–22 °C, under the PPFD of 40 µmol m^−2^ s^−1^ (12 h light/12 h dark), in 35 ppt artificial seawater (Red Sea salt, Red Sea Europe) and fed with *A. acetabulum* approximately once a week, as described in detail by Havurinne & Tyystjärvi (2020).

### Observations of cytoplasmic streaming with light microscopy

Algae were gently placed in a well of a 24 well-plate containing 1 mL of artificial seawater, after which light microscopy imaging was conducted with an inverted microscope (DMi1; Leica Microsystems, Germany), equipped with an HI Plan I 20x/0.3 PH1 objective (Leica). 10 μL of cytochalasin B (final concentration 100 μM) or oryzalin (final concentration 10 μM), for *A. acetabulum* or *Bryopsis* sp., respectively, was added, and imaging was continued after 20 min acclimation time. The microscope light was kept on during the whole measurement period. Stock solutions of cytochalasin B and oryzalin were prepared in dimethyl sulfoxide (DMSO). The experiments were repeated with 3 individual algal cells for both species.

### Macro-photography of chloroplast movements

Algae were placed in a 24 well-plate containing 1 mL artificial seawater and allowed to acclimate for 20–60 min under low light (PPFD ∼5–20 µmol m^−2^ s^−1^) at room temperature. In the case of 100 µM cytochalasin B (*A. acetabulum* only), algae were let to acclimate for further 20 min after the addition. Two types of experiments were conducted: cutting (both algae) and high light illumination (*A. acetabulum* only). In both cases, the well-plate was placed on a temperature-controlled box (ColdPlate; Qinstruments, Germany), set to 20 °C. In the case of cutting, an individual algal cell was first cut by scissors (avoiding touching any other cells in the well) into 2 pieces and the loose part (tip) was removed. In a separate experiment, algae were cut into pieces (*A. acetabulum* only) and 200 µL of the surrounding media (or clean media, as a mock treatment) was immediately transferred to another well with intact algal cells. After the cutting or addition of the media, the cells were kept on the temperature-controlled box under low light (PPFD ∼5–20 µmol m^−2^ s^−1^) for 30 min, after which a replicate (*A. acetabulum* only) was let to recover overnight under low light at room temperature (∼20 °C). In the case of the high light illumination, algae were exposed to light (PPFD of ∼1000 µmol m^−2^ s^−1^) from a 50W white LED flood light (ExtraStar Electrical Ltd; for the spectra, see Fig. S1) for 15 min, followed by 15 min of darkness. During both treatments (cutting and illumination), photographs were taken every 1 to 10 min with a Canon EOS 7D MKII camera (Canon, Japan), equipped with a Laowa 100mm f2.8 Ultra Macro APO 2:1 macro-objective (Laowa, China) and a speedlight flash. Settings were ISO 100, f4.5 and 1/200s exposure time. The experiments were repeated 2 to 4 times; each time, at least 3 different algal cells were placed in the well.

In the case of high light illumination and cutting/media addition with *A. acetabulum*, RGB (red, green and blue) images recorded with the Canon EOS 7D MKII were analysed with Fiji (ImageJ v1.54f; Schindelin 2012); 2 to 4 areas of interest (AOI; a well-focused algal segment with a length of a few mm), each one from an individual alga, were selected from each image (Fig. S2). Heterogeneous segments were avoided. For each selected AOI, mean intensities of the RGB channels were extracted using Fiji’s Colour Histogram plug-in, which analyses each channel independently on an 8-bit grayscale (0–255). The blue RGB channel provided the highest contrast between algal cells and the background and was therefore used to estimate chloroplast abundance within the selected algal segments; where lower blue-channel intensity corresponded to higher chloroplast abundance. Additionally, the standard deviation of blue-channel intensities within each selected AOI was quantified using the Colour Histogram plug-in to estimate the heterogeneity of chloroplast distribution. To assess whether scattered light affected the quantification, an empty background region containing only artificial seawater was additionally selected and analysed in each image. Average blue-channel intensities of the background showed only minimal variation throughout the experiment (Fig. S3).

### Imaging of herbivory by sea slugs with light microscopy

*A. acetabulum* cells were placed in a 24 well-plate containing 1 mL artificial seawater and allowed to acclimate for 20–60 min. The algae were then imaged at room temperature (∼20 °C) under low light (PPFD ∼5–20 µmol m^−2^ s^−1^) using a digital measuring microscope (DMS300; Leica) equipped with a 0.8x objective. A small (<10 mm in length) *E. timida* sea slug individual was then added to the well, and imaging was continued every 1–25 minutes, depending on the feeding activity of the slug, for 30–50 min before the sea slug was removed. After feeding, 2 mL artificial seawater was added to compensate for evaporation, and the algae were allowed to recover for 3 days under low light at room temperature. The experiment was repeated 3 times with 3 individual sea slugs and at least 3 individual algal cells per experiment.

### Chlorophyll *a* fluorescence

Pulse amplitude modulated (PAM) chlorophyll *a* fluorescence was recorded with the Mini version of Imaging-PAM fluorometer (Walz, Germany). Algae were placed in a 24 well-plate containing 1 mL artificial seawater and dark-acclimated for 20 min in the absence or presence of 100 µM cytochalasin B for *A. acetabulum* or 10 µM oryzalin for *Bryopsis* sp.. Maximum PSII activity was estimated with F_V_/F_M_, calculated as (F_M_-F_O_)/F_M_ and measured using low-intensity measuring flashes (setting 2; frequency 1 Hz) and a saturating light pulse (setting 9; 600 ms), where F_O_ and F_M_ represent the minimum and maximum chlorophyll *a* fluorescence yields of dark-acclimated samples, respectively. Algae were then exposed to three different blue light treatments (for the spectra, see Mattila et al. 2025): (1) 15 min high light (PPFD ∼1000 µmol m^−2^ s^−1^) followed by 15 min of darkness, 15 min high light and 20 min darkness (Figs 1A–D and 2 A–D), (2) the same high light/darkness illumination, but half of the algae were darkened with a black, 3D-printed plastic cover (designed with FreeCAD and printed with Original Prusa MK4; Prusa Research, Czechia; Fig. S4) to create a sharp light gradient across the algal sample (Figs 1E–H and 2 E–H), and (3) 5 min of gradually increasing light (PPFD from ∼0 to 1000 µmol m^−2^ s^−1^) followed by 5 min with gradually decreasing light (PPFD from ∼1000 to 0 µmol m^−2^ s^−1^), repeated for five times (for more details, see Mattila et al. 2025) and finally, by 20 min of darkness (Fig. S5). During all the above-described light treatments, saturating light pulses were fired every 60 s (unless otherwise described) to calculate NPQ, as (F_M_-F_M_’)/F_M_’, and PSII “operational efficiency”, as (F_M_’-F’)/F_M_’, where F_M_’ is the maximum (during a saturating light pulse) and F’ the incident chlorophyll *a* fluorescence yield under illumination. For analysis of fluorescence parameters, 3 areas of interest were selected from each analysed algal cell. In the case of the “half-shade” experiment (Figs S4 and S5), the fluorescence parameters from shaded parts were quantified using saturating light pulses before placing the cover (before switching on the light) and at the end of the treatment, after removing the cover. The experiments were repeated 2 times, each time with 3 to 4 different algal cells, for both species.

### Figures and statistical analyses

Statistically significant differences in photoinhibition of PSII were tested using the Mann-Whitney U test with Microsoft Excel using the Real Statistics Resource Pack (Zaiontz 2020). Macro-imaging time-course data were analysed with a Linear Mixed Effects Model (LMM) using the lme4 package in R v4.6.0 (Bates et al. 2015; R Core Team 2021), where treatment and time were assigned as fixed effects and individual replicate as a random effect. Homoscedasticity and LMM fit were evaluated by visual inspection of the residuals vs. fitted values plots, and normality of residuals was evaluated by inspecting Q-Q plots and histograms of residual distributions. The LMM was followed by a Type III ANOVA with Satterthwaite’s approximation for degrees of freedom using the lmerTest package (Kuznetsova et al. 2017) to test the effects of treatment, time and the interaction of treatment x time. Post hoc pairwise or multiple comparisons of estimated marginal means were done with the emmeans package (Lenth & Piaskowski 2026). Bonferroni-adjusted *p*-values were used in multiple comparisons. Figures were prepared in R with the ggplot2 package (Wickham 2016).

## Results

### *A. acetabulum* showed more rapid cytoskeleton-controlled chloroplast movements than *Bryopsis* sp

Bidirectional chloroplast movements, along the cell’s longitudinal axis, were observed in *A. acetabulum* and *Bryopsis* sp. under light microscope (Fig. 1). Based on time-series of the microscopy images, the chloroplast movement rates were calculated to be approximately 149 ± 52 and 30 ± 15 µm min^−1^, for *A. acetabulum* and *Bryopsis* sp., respectively. Treatments with cytochalasin B (an inhibitor of actin polymerization) in *A. acetabulum* or with oryzalin (an inhibitor of microtubule assembly) in *Bryopsis* sp. strongly inhibited, and in many cases almost completely abolished, the chloroplast movements (Fig. 1), in agreement with previous literature (e.g., Nagai & Fukui 1981; Ochiai et al. 2024). Thus, the results indicate that chloroplast movements in these species were indeed controlled by the cytoskeleton.

### Prevention of chloroplast movements increased PSII photoinhibition in *A. acetabulum* but not in *Bryopsis* sp

To test whether the chloroplast movements respond to high light and constitute a photoprotective mechanism, chlorophyll *a* fluorescence kinetics under light were measured from *A. acetabulum* (Fig. 2) and *Bryopsis* sp. (Fig. 3) in the presence and absence of the movement inhibitors cytochalasin B and oryzalin. First, dark-acclimated algae were exposed to alternating 15 min periods of high light (PPFD of 1000 µmol m^−2^ s^−1^) and darkness (Figs 2A–D and 3A–D). Then, to test if chloroplast movements would be enhanced in the presence of a partial shade, the treatment was repeated while covering half of the algal sample (Figs 2E–H and 3E–H; for the cover, see Fig. S1).

**Figure 2.**
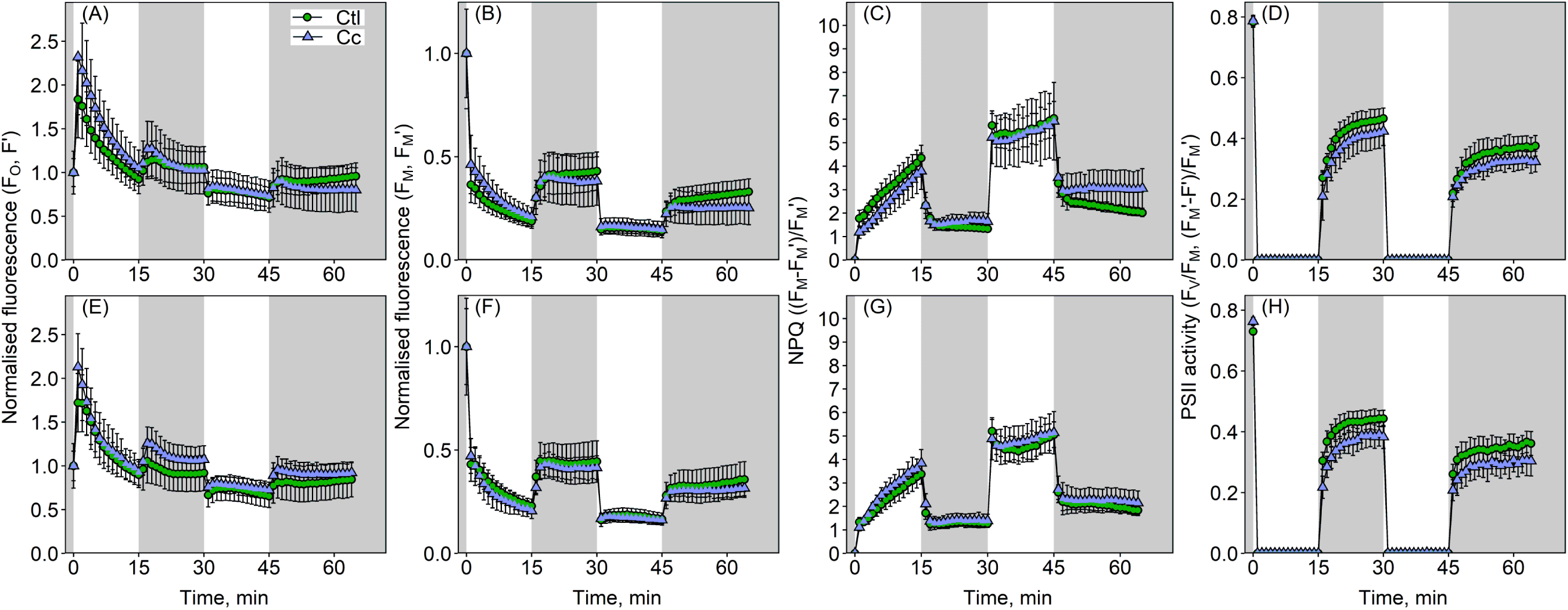
Effects of the inhibition of chloroplast movements with cytochalasin B on chlorophyll *a* fluorescence kinetics in *Acetabularia acetabulum*. Before the treatments, algae were dark-acclimated for 20 min, in the absence (Ctl; circles) or presence of cytochalasin B (Cc; triangles). In (A–H), algae were treated with periods of high light (PPFD 1000 µmol m^−2^ s^−1^; white background) and darkness (grey background). In (E–H), half of the algae were shaded (see Fig. S1); the measurements shown are from the illuminated parts. (A and E) Minimum fluorescence of dark-acclimated samples (F_O_) or incident fluorescence under illumination (F’), normalised to the starting values. (B and F) Maximum fluorescence, obtained by firing saturating light pulses, of dark-acclimated samples (F_M_) or under illumination (F_M_’), normalised to the starting values. (C and G) Non-photochemical quenching (NPQ) and (D and H) F_V_/F_M_ (dark-acclimated sample) or PSII operational yield (under illumination). The symbols show averages based on six to eight biological replicates and the error bars show standard deviations.

**Figure 3.**
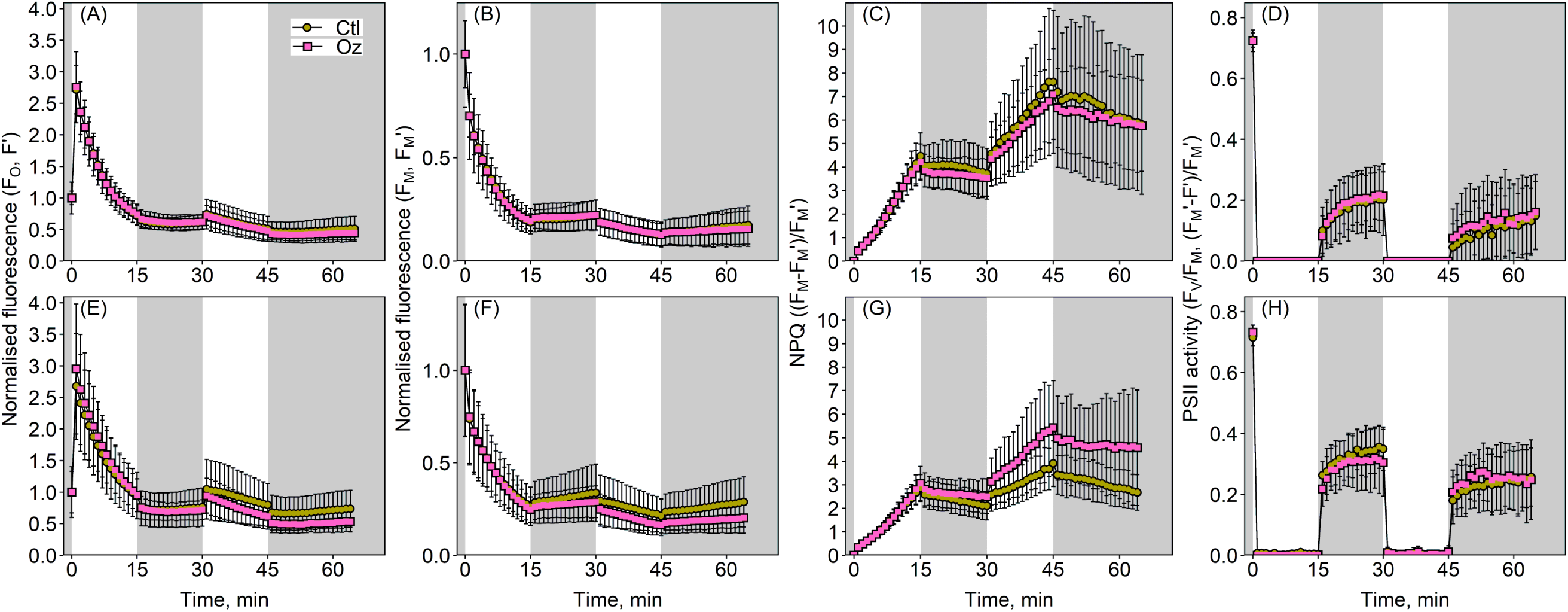
Effects of the inhibition of chloroplast movements with oryzalin on chlorophyll *a* fluorescence kinetics in *Bryopsis* sp.. Before the treatments, algae were dark-acclimated for 20 min, in the absence (Ctl; circles) or presence of oryzalin (Oz; squares). In (A–H), algae were treated with periods of high light (PPFD 1000 µmol m^−2^ s^−1^; white background) and darkness (grey background). In (E–H), half of the algae were shaded (see Fig. S1); the measurements shown are from the illuminated parts. (A and E) Minimum fluorescence of dark-acclimated samples (F_O_) or incident fluorescence under illumination (F’), normalised to the starting values. (B and F) Maximum fluorescence, obtained by firing saturating light pulses, of dark-acclimated samples (F_M_) or under illumination (F_M_’), normalised to the starting values. (C and G) Non-photochemical quenching (NPQ). (D and H) F_V_/F_M_ (dark-acclimated sample) or PSII operational yield (under illumination). The symbols show averages based on six to eight biological replicates and the error bars show standard deviations.

If chloroplasts move away from high light, chlorophyll *a* fluorescence should decrease and if they return during a subsequent darkness, fluorescence should again increase, and the use of the cytoskeleton inhibitors should allow the differentiation of these effects from other changes (high) light induces on the fluorescence kinetics (such as NPQ). However, inhibition of chloroplast movements did not substantially affect fluorescence kinetics in either species; consequently, NPQ and PSII operational yield were also minimally affected (Figs 2 and 3). Regardless, in *A. acetabulum*, incident and maximum fluorescence in the light (F’ and F_M_’) were slightly higher at the beginning of the illumination and often lower during dark in the presence of cytochalasin B, as compared to control treatments without the inhibitor (Fig. 2A, B, E and F). Consequently, cytochalasin B reduced NPQ at the onset of the illumination and increased NPQ during darkness; however, this effect was not observed in the shading treatment (Fig. 2C and G). PSII operational efficiency (F_M_’-F’)/F_M_’ was consistently lower in the presence of cytochalasin B (Fig. 2D and H). PSII photoinhibition, quantified as the percentage of the initial F_V_/F_M_ values at the final time-point, increased in the presence of cytochalasin B; the difference was statistically significant both without and with the half-shade cover (Table 1; *p* = 0.010 and *p* = 0.031, respectively, according to Mann-Whitney tests). On the other hand, the F_V_/F_M_ values quantified from the shaded regions of the algae were similar in the absence and presence of cytochalasin B (Table 1), suggesting that cytochalasin B did not induce inactivation of PSII in the dark.

**Table 1.** Quantification of PSII photoinhibition. PSII activity, probed by the FV/FM chlorophyll *a* fluorescence parameter, as a percentage of the initial values (± SD), after high light treatments in *Acetabularia acetabulum* and *Bryopsis* sp. in the absence (−) or presence (+) of inhibitors of chloroplast movements (cytochalasin B for *A. acetabulum* and oryzalin for *Bryopsis* sp.). The treatments involved periods of high light and darkness (on/off fluctuating light (FL); Figs 2 and 3), half of the algae shaded when indicated (Half-shade; values from illuminated parts of the algae, Figs 2 and 3), or periods of gradually increasing and decreasing light (Fig. S5). The values in the last column (Dark) have been quantified from algae kept under the “half-shade” condition (Fig. S4), after removing the cover. Statistically significant differences in PSII activity at the *p* < 0.05 level, compared to non-inhibitor treated samples, are highlighted with bold text and asterisks.

| Species | PSII activity, % of initial $F_v/F_m$ | | | | |
| --- | --- | --- | --- | --- | --- |
|  | Inhibitor | On/off FL | Half-shade | Gradual FL | Dark |
| *A. acetabulum* | - | **47.9 ± 3.9** | **51.3 ± 5.0** | 65.8 ± 3.8 | 75.1 ± 14.1 |
| *A. acetabulum* | + | **41.1 ± 3.4\*** | **43.2 ± 5.7\*** | 60.7 ± 11.6 | 75.5 ± 6.2 |
| *Bryopsis* sp. | - | 20.6 ± 14.9 | 38.6 ± 14.7 | 42.5 ± 6.2 | 65.0 ± 20.9 |
| *Bryopsis* sp. | + | 21.7 ± 15.7 | 34.5 ± 18.7 | 36.9 ± 10.1 | 50.1 ± 32.1 |

In *Bryopsis* sp., oryzalin did not affect chlorophyll fluorescence yields (F’ and F_M_’) in the first 15 min light/15 min dark treatment (Fig. 3A and B). In contrast, oryzalin decreased both F’ and F_M_’ under the partly shaded light treatment, but without clear light/dark dependence (Fig. 3E and F). In the latter treatment, NPQ increased in the presence of oryzalin (Fig. 3G). No statistically significant effects on PSII photoinhibition were found at the final time-points in either of the treatments (Table 1).

Finally, we also tested if changing the fluctuating light treatment (this time, light intensity gradually increased for 5 min from PPFD 0 to 1000 µmol m^−2^ s^−1^, and then decreased for 5 min; the cycle was repeated 5 times) would affect the results (Fig S5). In *A. acetabulum*, the effects of cytochalasin B on fluorescence yields and parameters were similar to those shown in Fig. 2, albeit less clear, and therefore the effect on PSII photoinhibition was not statistically significant (Fig. S5A–D; Table 1). In *Bryopsis* sp., PSII operational efficiency was slightly lower in the presence of oryzalin than in its absence (Fig. S5E–H), but again, the difference was not statistically significant (Table 1).

### Reversible chloroplast aggregation was observed in *A. acetabulum* in response to a high light treatment

To confirm that the differences in chlorophyll *a* fluorescence kinetics in *A. acetabulum* in the absence and presence of cytochalasin B (Fig. 2) were related to chloroplast movements, and to further characterize these movements, *A. acetabulum* was again subjected to a treatment with 15 min high light (PPFD 1000 µmol m^−2^ s^−1^)/15 min darkness, in the absence or presence of cytochalasin B, during which macro-imaging was performed (Fig. 4). To get a quantitative estimate of chloroplast density, average pixel intensities in the blue channel of selected algal segments were quantified from the captured RGB-images (Figs 4A and S2; for the selection criteria, see Materials and methods). In addition, evenness of the chloroplast distribution in each selected algal segment was estimated by quantifying standard deviation (in the blue channel values) within the selected areas of interest (Fig. 4B). In the absence of the movement inhibitor cytochalasin B, pixel intensities (in the blue channel) increased during the high light period, especially at the beginning of the illumination, and decreased during the subsequent darkness (Fig. 4A), suggesting that the average chloroplast density in the selected algal segments was transiently diminished during the high light illumination. Variation in the pixel values (in the blue channel) of the selected algal segments also increased during the high light period and, during the subsequent darkness, decreased back to the original levels (Fig. 4B), suggesting that chloroplasts were distributed less homogeneously under high light than under darkness. The treatment (presence or absence of cytochalasin B), time and the interaction between the treatment and time had statistically significant effects on both the mean pixel values and the variation (Type III ANOVA from LMM, *p* < 0.05; Tables S1 and S2), suggesting that cytochalasin B substantially suppressed the above-described light-induced responses in chloroplast distribution (see Fig. 4A and 4B for post hoc pairwise comparisons throughout the whole time course of the experiment). Indeed, visual examination of the RGB-images revealed small dark dots, likely reflecting chloroplast aggregations, under high light, which largely disappeared upon the subsequent darkness (Fig. 4C–E).

**Figure 4.**
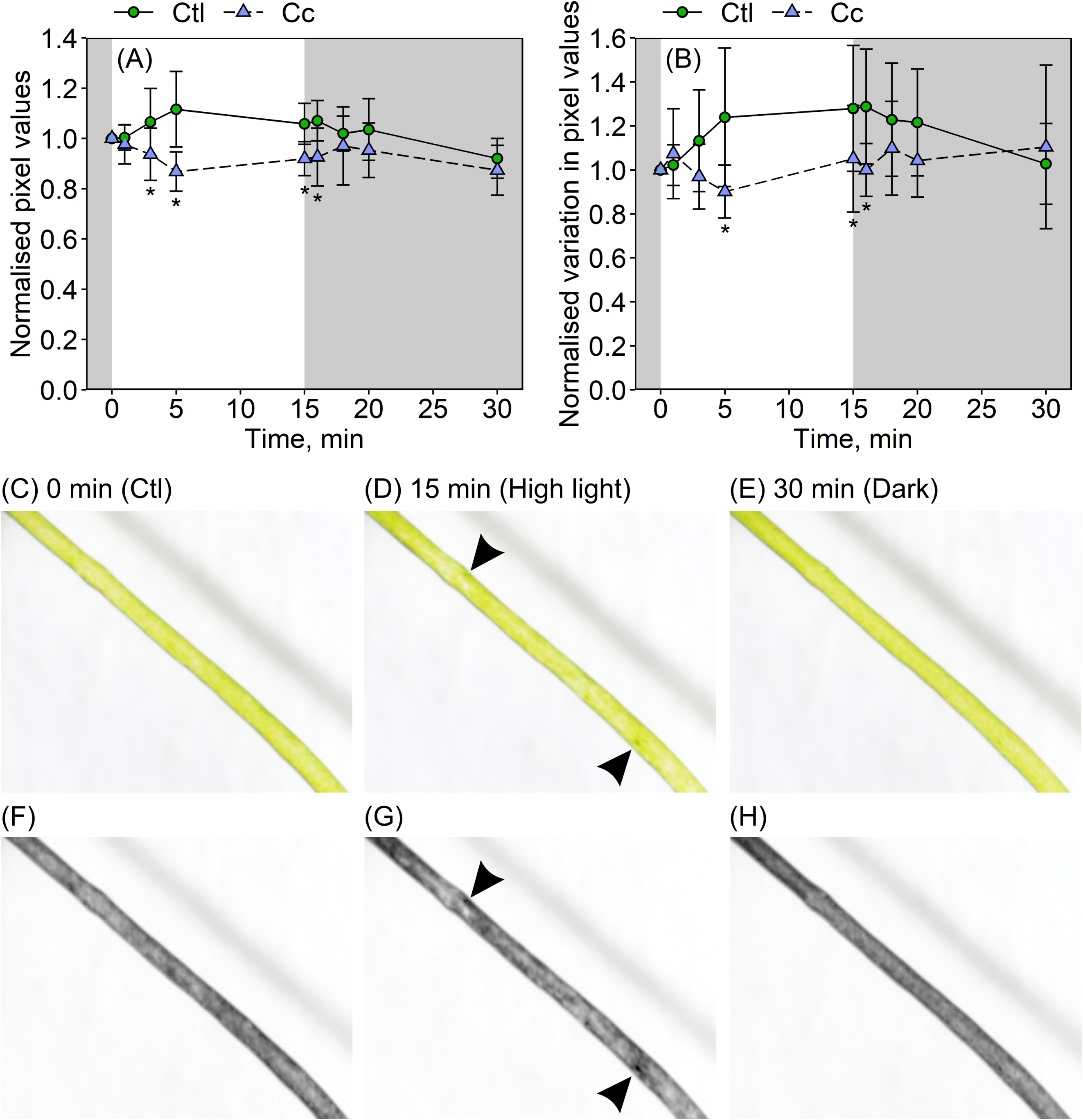
Macro-imaging revealed reversible chloroplast aggregation in *Acetabularia acetabulum* cells during a high light treatment. Low-light-acclimated algae were illuminated for 15 min with high light (PPFD ∼1000 µmol m^−2^ s^−1^) and subsequently incubated in darkness for 15 min, in the absence (Ctl; circles) or presence of cytochalasin B (Cc; triangles). (A) Average values, reflecting overall chloroplast density (higher values reflecting lower densities), and (B) standard deviations, reflecting spatial heterogeneity in chloroplast distribution (higher values reflecting greater heterogeneity, perhaps chloroplast aggregation), in the blue channel values within the regions of interest, quantified from selected algal segments (Fig. S2) from the captured RGB-images and normalised to their starting values. Symbols show averages and error bars standard deviations based on 12 biological replicates. Asterisks indicate statistically significant differences (*p* < 0.05*) between the treatments at individual time points (according to linear mixed model, with post hoc estimated marginal means pairwise comparisons). (C–H) An exemplary image series showing a part of an algal cell at the start of the treatment (0 min), after 15 min of high light illumination and after subsequent 15 min in darkness, in the absence of cytochalasin B. (C–E) Original RGB-images and (F–H) the blue channel (used for the quantifications), represented as grey-scale images. The arrowheads highlight some of the darker dots (presumably aggregated chloroplasts) that appeared during the high light illumination and disappeared during the subsequent darkness.

### Wounding led to systemic chloroplast aggregation in *A. acetabulum* but not in *Bryopsis* sp

Although chloroplast movements (aggregation of chloroplasts, in particular) were detected in response to high light, as described above, they were of limited magnitude. *A. acetabulum* is known to be sensitive to handling (Fester et al. 1994) and, therefore, we next examined the responses of *A. acetabulum* and *Bryopsis* sp. cells to mechanical damage, using the macro-imaging approach. After an *A. acetabulum* cell was cut into two pieces with scissors, the appearance of darker and lighter regions (presumably due to chloroplast aggregation) could be visually detected in the cut cell within 1–2 min and uncut neighbour cells were also similarly affected (Fig. S6). Accordingly, a clear increase in the blue channel pixel values of selected uncut cells was observed, especially 10–30 min after the cutting (Fig. 5A), possibly due to increased non-green segments (i.e., chloroplast-empty algal regions) and/or decreased average number of chloroplasts in the quantified segments. Variation in the selected algal segments (in the blue channel) also strongly increased after the cutting (Fig. 5B), suggesting that the cutting triggered strong chloroplast aggregation. The aggregations were not permanent, as cells allowed to recover overnight regained a uniform green colour (Fig. 5C). To confirm that the observed chloroplast aggregation was an active response controlled by the cytoskeleton, the experiment was repeated with the actin inhibitor cytochalasin B (Fig. S7). In this case, some of the used algae showed inhomogeneous chloroplast distribution already before cutting; regardless, cutting further increased chloroplast aggregation only in the absence of cytochalasin B.

**Figure 5.**
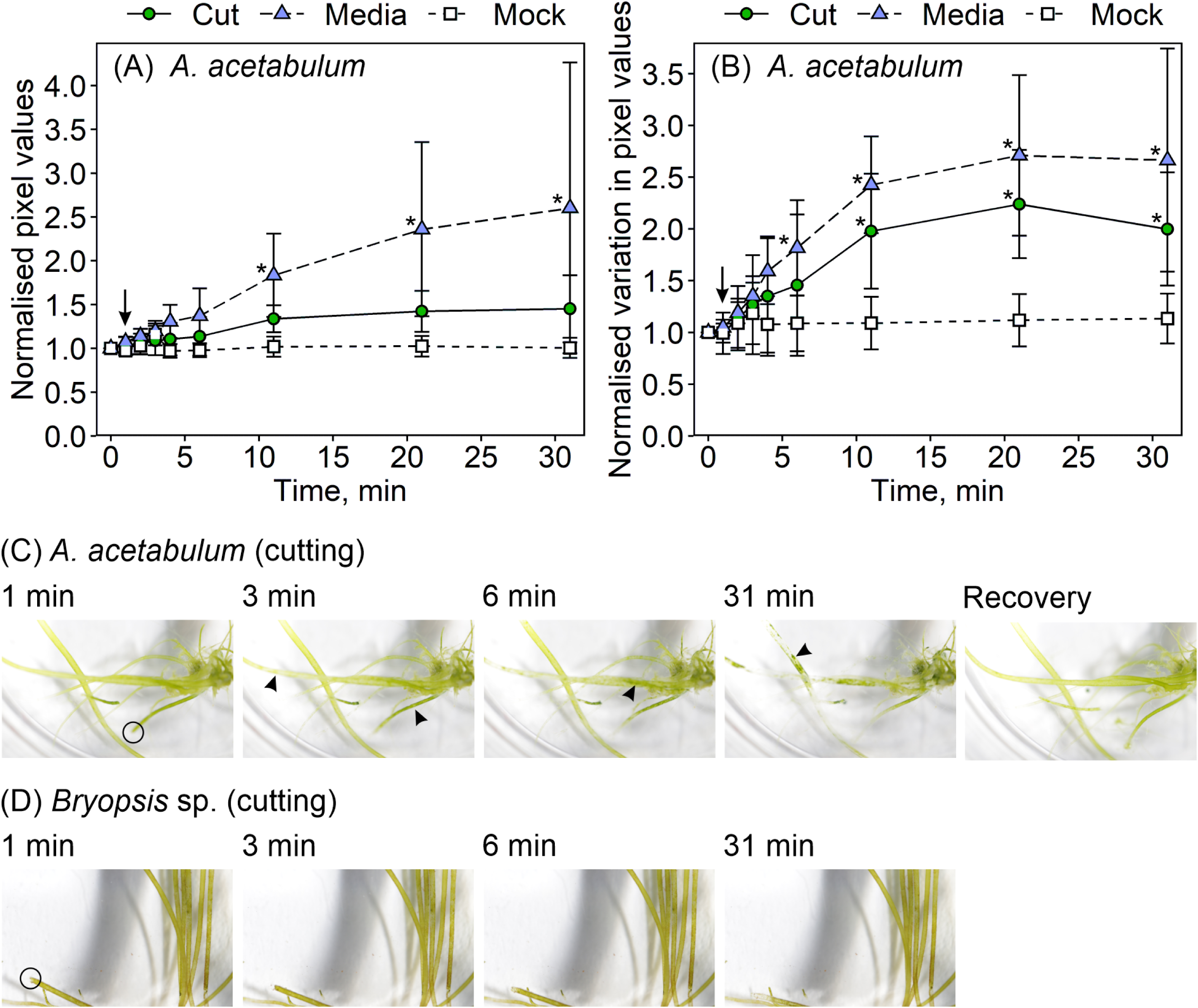
Chloroplasts aggregated in *Acetabularia acetabulum*, but not in *Bryopsis* sp., in response to wounding, as observed with macro-imaging. A low-light-acclimated cell of *A. acetabulum* (A–C) or *Bryopsis* sp. (D) was cut with scissors (Cut), or 200 µL of artificial seawater medium, obtained from another well where *A. acetabulum* cells were cut into pieces right before the collection of the medium, was added to the well with intact algae (Media) or 200 µL of clean artificial seawater medium was added to the well (Mock). The timepoints highlighted with the arrows (A and B) represent an image taken immediately after the cutting or media additions. Algae were then incubated under low light up to 30 min, and (A) average values, reflecting overall chloroplast density (higher values reflecting lower densities), and (B) standard deviations, reflecting spatial heterogeneity in chloroplast distribution (higher values reflecting greater heterogeneity, perhaps chloroplast aggregation), in the blue channel values within the regions of interest, quantified from selected algal segments (Fig. S2) from the captured RGB-images and normalised to their starting values. Symbols show averages, and error bars show standard deviations based on 4–6 biological replicates. Asterisks highlight statistically significant differences (*p* < 0.05*) between the cutting or addition of media treatments and the mock treatment at the indicated time points (according to linear mixed model, with post hoc estimated marginal means multiple comparisons using Bonferroni adjusted *p-*values). (C and D) An exemplary image series showing algal cells right after the cutting (1 min) and after further 2, 5 and 30 min, and in the case of (C), after an overnight recovery, as indicated. The black circles highlight the cut cells (first images). The arrowheads highlight the first appearances of darker and/or lighter dots, presumably indicating chloroplast aggregation. For the whole time-series and more biological replicates, see Fig. S6.

To test whether compounds released from cut *A. acetabulum* cells into the surrounding media can induce chloroplast aggregation in uncut cells, a cell was cut to pieces in a separate well-plate. A small amount of the medium was then immediately transferred to another well-plate containing intact algae (Figs 5A, 5B and S6). As with the cutting, both the average pixel values and variation (in the blue channel) clearly increased after the addition of the medium (Figs 5A and 5B). In contrast, a mock treatment (addition of clean artificial seawater medium) did not alter chloroplast density or distribution, as probed with the macro-imaging (Figs 5A and 5B and S6).

When all the treatments (cutting, addition of media from wounded algae and mock) were taken into account, the treatment, time as well as the interaction effect of treatment and time were statistically significant on both the mean pixel values and the standard deviations (Type III ANOVA from LMM, *p* < 0.05; Tables S3 and S4). Post hoc analyses revealed that, compared to the mock treatment with clean artificial seawater, only the addition of medium from wounded algae had a significant effect on the chloroplast density, when probed by mean pixel values, whereas both the addition of medium from wounded algae and the cutting treatment responses differed statistically significantly from the mock treatment (see Fig. 5A and 5B for the pairwise comparisons throughout the whole experimental time course).

Unlike *A. acetabulum*, *Bryopsis* sp. cells responded to cutting primarily locally, i.e., chloroplast aggregation was observed almost exclusively in the cut cells, and close to the cutting site (Figs 5D and S6).

In a natural context, wounding may derive from herbivory. Therefore, the sea slug *E. timida*, a natural herbivore of *A. acetabulum*, was placed in a well-plate containing intact algae (Figs 6 and S8). *E. timida* feeding on an alga eventually drained the cell of most of its chloroplasts and the algal cell became nearly transparent. Similarly to the cutting experiment (Fig. 5C), signs of chloroplast aggregation could be visually detected already within the first 1–2 min of the feeding in the algal cells not touched or damaged by the sea slug (as judged by visual inspection) (Fig. 6). After 10–30 min of feeding, cells distant from the sea slug (feeding site) also showed chloroplast aggregation. After the sea slug was removed and the alga was left to recover for 3 days under low light, all cells recovered, except the cell which had been fed on (Fig. S9).

**Figure 6.**
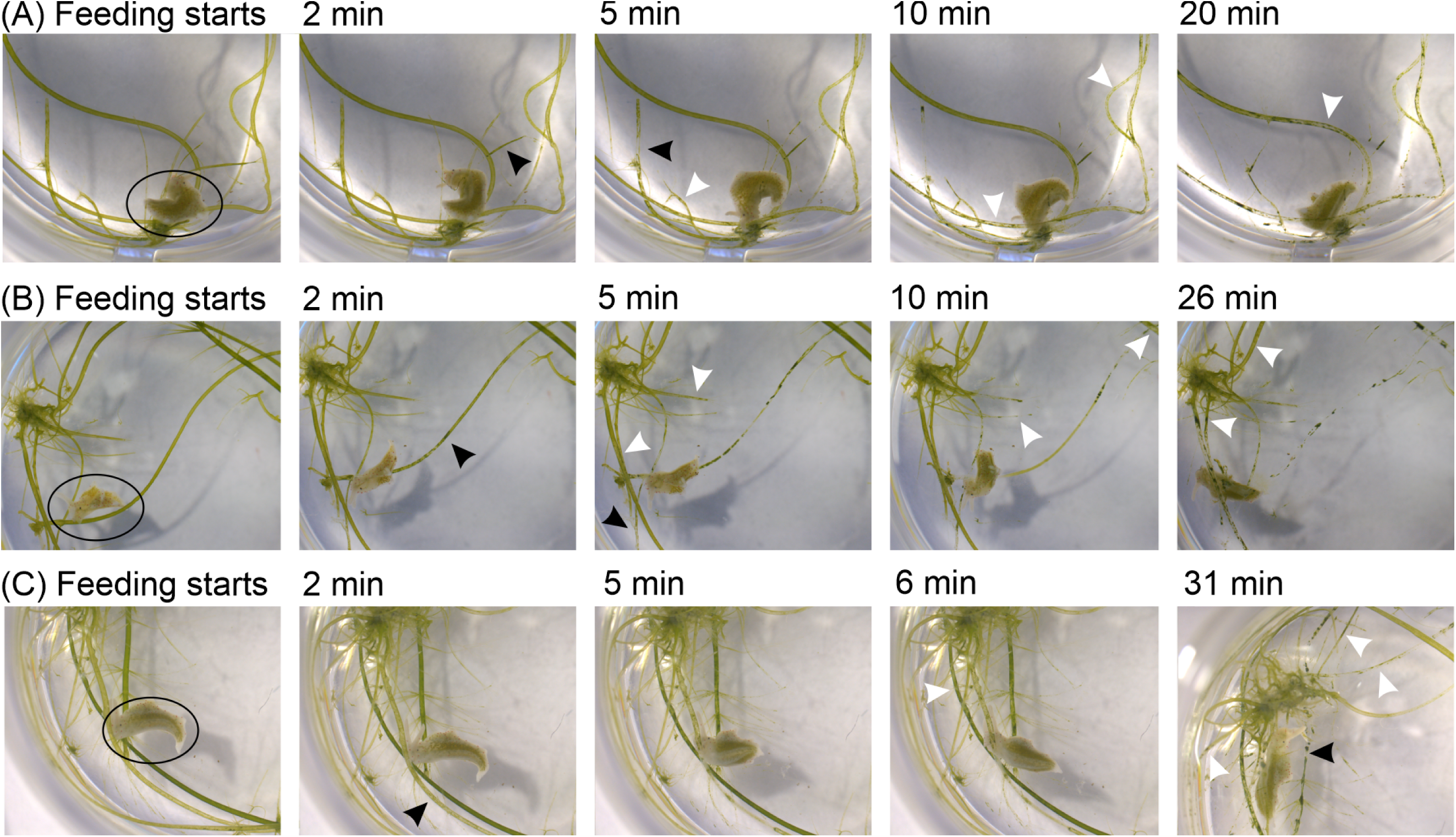
Chloroplast aggregation in *Acetabularia acetabulum* cells in response to grazing by an *Elysia timida* sea slug, as followed with digital measuring microscope. (A–C) 3 biological replicates. A sea slug (highlighted with a circle in the first image of each series) was placed in a well with intact algal cells; the indicated time-points show how much time has passed since the sea slug began feeding on an algal cell. Black arrows highlight chloroplast aggregation (appearance of lighter/darker spots) within the algal cell the sea slug is feeding on, presumably caused by the sucking of the slug. The white arrows highlight chloroplast aggregation in algal cells, on which the sea slug has not fed (based on visual inspection). Brightness and contrast have been adjusted equally in all the images in each series. For the full time-series and original images, see Fig. S8.

## Discussion

The two green siphonous macroalgae studied here showed different responses to abiotic and biotic stressors. The Dasycladalean alga *A. acetabulum* responded to high light with reversible aggregation of chloroplasts (Figs 2 and 4) and cellular damage, caused by cutting or feeding by the sacoglossan sea slug *E. timida*, induced even stronger chloroplast aggregations (Figs 5 and 6). In contrast, such responses were not observed in the Bryopsidales macroalga *Bryopsis* sp. (Figs 3 and 5). Chloroplasts in *Bryopsis* sp. have been previously shown to move at the rates of 20–60 µm min^−1^ (Menzel & Schliwa 1986; Ochiai et al. 2024). A similar rate (∼30 µm min^−1^) was measured also in the present study. Thus, the lack of observed chloroplast movements was not due to an inability of the Bryopsidales alga for rapid cytoplasmic streaming. However, we observed even faster chloroplast movements in *A. acetabulum* (∼150 µm min^−1^), perhaps in line with the readiness of the alga to respond to the tested stresses with the movements (most notably, via the chloroplast aggregation).

### Is chloroplast aggregation a photoprotective response?

High-light-induced chloroplast aggregation, as observed here for *A. acetabulum* (Fig. 4), has been reported in phylogenetically distant algae, including large diatom species (Kiefer 1973; Silkin et al. 2021) and the siphonous yellow-green macroalga *V. sessilis* (Blatt 1983). In *A. acetabulum*, chloroplasts have been previously shown to accumulate in the stalk under low/moderate light and move away from the stalk during darkness (night) (Schmid & Koop 1983; Paques & Brouers 1984), but high-light-induced responses have not previously been described. During illumination, but not in darkness, inhibition of chloroplast movements by cytochalasin B (Fig. 1) caused a small but consistent decrease in the PSII capacity, as indicated by reduced F_V_/F_M_ values (Fig. 2; Table 1). This suggests that light avoidance movements and/or chloroplast aggregation (Fig. 4) protect PSII from photoinhibition in *A. acetabulum*, as previously reported for plants and diatoms (e.g., Kasahara et al. 2002; Silkin et al. 2021; Bastos et al. 2025).

Aggregation might protect chloroplasts by enhancing shading. In the unicellular diatom *Pleurosigma strigosum*, PSII activity (probed by the F_V_/F_M_ parameter) recovers faster and less NPQ is induced in the central part of the contracted chloroplast, compared to the distal parts (Bastos et al. 2025). On the other hand, in some large centric diatoms, chloroplast aggregation is not related to photoprotection but reflects cell death (Goessling et al. 2016). Additionally, the amount of self-shading is expected to be low in optically thin samples, such as in a single microalgal cell and, to some degree, also in individual macroalgal filaments, such as the *A. acetabulum* cells used in the present study. In plants, besides the self-shading, chloroplast movements may protect chloroplasts by avoiding extremely bright spots, which are created due to a lens-effect of epidermal cells (Gan et al. 2025).

The photoprotective role of chloroplast movements has also been debated in plants, mainly because most of the studies have relied on *in vivo* chlorophyll *a* fluorescence measurements, which themselves are affected by chloroplast movements (Wilson & Ruban 2020), although compared to plants, the less complex optical properties of the tubular cells of *A. acetabulum* and *Bryopsis* sp. might make fluorescence measurements less complicated (Gan et al. 2025). Nevertheless, here PSII photoinhibition was assessed solely by chlorophyll *a* fluorescence; therefore, additional approaches, such as oxygen evolution or PSI activity, are needed to verify the photoprotective role of chloroplast movements in *A. acetabulum*, particularly under natural conditions.

We hypothesised that *Bryopsis* sp. employs chloroplast movements as a compensatory photoprotective mechanism, as the alga lacks several common photoprotection mechanisms (e.g., Christa et al. 2017; Iha et al. 2021; Havurinne et al. 2025). In *A. acetabulum*, incident and maximum yields of chlorophyll *a* fluorescence (F’ and F_M_’) were usually lower in the absence than in the presence of a movement inhibitor, especially at the start of the high light illumination, and often also increased more rapidly in the darkness (Fig. 2). This pattern is consistent with high-light-induced chloroplast avoidance and accumulation responses, respectively. However, inhibition of chloroplast movements in *Bryopsis* sp. with oryzalin did not cause changes in the chlorophyll *a* fluorescence kinetics that would be consistent with light-induced chloroplast movements (Fig. 3). Oryzalin also did not increase PSII photoinhibition (Table 1). Instead of chloroplast movements, Bryopsidales algae might rely on shading caused by whole filaments (protecting the lower filaments of the algal tuft), sustained NPQ, accumulation of certain carotenoids and/or oxygen-dependent auxiliary electron transfer pathways (e.g., Giovagnetti et al. 2018; Giossi et al. 2021; Mattila et al. 2025). Recently, we observed that *Bryopsis* sp. repairs PSII damage slowly, as compared to *A. acetabulum* (Mattila et al. 2025). As both PSII repair and chloroplast movements require substantial amounts of energy, in the form of ATP (Murata & Nishiyama 2018; Wada & Kong 2018), *Bryopsis* sp. may instead tolerate high light stress rather than actively avoiding or repairing it, thereby conserving resources for other physiological processes.

In some Bryopsidales algae, peculiar chlorophyll *a* fluorescence kinetics have been observed, such as a strong decrease in the fluorescence yield (F’ or F_O_’) during illumination, well below the original F_O_ level (e.g., Fig 3; Mattila et al. 2025), an increase from F_M_ to F_M_’ under low or moderate light (Havurinne et al 2025) and a decrease in F_M_’ shortly after switching off a high light illumination (Seki et al. 2025). As no high-light-induced chloroplast movements were observed in *Bryopsis* sp., other explanations will be needed to explain these observations.

### Is chloroplast aggregation a defence strategy against herbivory?

Some algal herbivores, including kleptoplastidic sea slugs, pierce the algal cell wall to suck out the cytoplasm, including chloroplasts. In *A. acetabulum*, both cutting and feeding by the “sap-sucking” sea slug *E. timida* caused strong chloroplast aggregation within minutes of the damage (Figs 5 and 6). These effects were observed not only at the damage site but also in nearby undamaged cells. Therefore, *A. acetabulum* might react to cellular damage as if by anticipating an herbivore attack. This might be beneficial since it would allow the alga to detect and prepare for a nearby herbivory attack, especially in the case of slowly moving herbivores, such as sea slugs and snails (see, Toth & Pavia 2000).

Herbivory can significantly reduce macroalgal growth, and in some cases, even determine species distribution (Duffy & Hay 2000; Poore et al. 2012; Friedlander & Critchley 2024; Terán et al. 2025). The sacoglossan sea slug *E. timida* is kleptoplastidic, i.e., it retains the “stolen” algal chloroplasts (kleptoplasts) functional for weeks, even months inside its animal body (de Negri & de Negri 1876; Cruz & Cartaxana 2022). The importance of these sea slugs in determining algal growth and distribution in nature is not well understood. Under laboratory conditions, sacoglossan species such as the non-kleptoplastidic *Aplysiopsis enteromorphae* and the kleptoplastidic *Elysia tuca*, greatly harm their prey algae *Chaetomorpha linum* and *Halimeda incrassata*, respectively (Trowbridge 1993; Rasher et al 2015). It has also been suggested that *Phyllaplysia taylori* controls the epiphytic algae growing on seagrasses (Hughes et al. 2010) and that local populations of the Bryopsidales algae *Codium* spp. are controlled by *Elysia viridis* and *Placida dendritica*, the latter of which cannot retain kleptoplasts functional (Trowbridge 1992; 2002). However, *Elysia subornata* could not control the growth of the invasive Bryopsidales alga *Caulerpa taxifolia* (Thibaut et al. 2001). Furthermore, in the field, contribution from the sap-sucking sea slugs to total herbivory was reported to be minor (Trowbridge 1993). Regardless, effects from the *E. timida* feeding are expected to be dramatic for an *A. acetabulum* individual, as the cell that the sea slug was feeding on showed no recovery even several days after removal of the slug (Fig. S9).

Algae have evolved several defence mechanisms against herbivory (for reviews, see Hansel & Diaz 2021; Burnett & Koehl 2022; Saha & Fink 2022; Lang et al. 2024), but to our knowledge, chloroplast aggregation has not previously been reported as a defence response to herbivory. *E. timida* is a specialist feeder of *A. acetabulum*, and thus, the alga could be expected to have developed specific defence strategies against it. For example, *H. incrassata* responds to the feeding by *E. tuca* by dropping the branches the sea slug is feeding on (Rasher et al. 2015). Does chloroplast aggregation make them harder for *E. timida* to acquire? Before starting to feed, *E. timida* usually probed the algae for some time (Fig. S6). In general, sea slugs rely on chemoreceptors to find their algal prey (Jensen 1982; Wertz et al. 2006; Bornancin et al. 2017). *Elysia* species have been shown to follow signals towards non-prey algae as well (Jensen 1988), suggesting that additional signals, such as components in the algal cell wall (Jensen 1997), are needed to select the algal prey. However, future research should determine if chloroplast aggregation hinders the feeding by *E. timida*. It should also be considered whether the chloroplast aggregation is the primary defence response. Possibly, the alga responds to an herbivore attack by massive withdrawal of all the major components of the cytoplasm, the chloroplast aggregation being just a side effect (or a part of the full response). Previous literature shows that *A. acetabulum* chloroplasts move away from the stalk to the base during the night (Schmid & Koop 1983; Paques & Brouers 1984). Some kleptoplastic sea slugs, e.g., *Plakobranchus ocellatus*, feed during nights (Tanamura & Hirose 2016) and thus, the above-described diurnal chloroplast movements could be an effort to try to avoid getting fed on, as empty *A. acetabulum* cells most probably are less attractive to the sea slug.

We observed that chloroplast aggregation in *A. acetabulum* spread from the primary damage site to distal areas (usually within 5 to 10 minutes) and even to undamaged cells (Figs 5 and 6), suggesting that water-borne signals or cues are involved in the process. Indeed, artificial seawater medium collected from a well with pre-cut cells was enough to induce chloroplast aggregation in intact cells (Fig. 5). Similarly, predation from the sea snail *Littorina obtusata* induced phlorotannin production (a defence mechanism) in grazed brown macroalgae *Ascophyllum nodosum*, as well as in ungrazed algae if introduced to water collected from the vicinity of the grazed algae (Toth & Pavia 2000). However, in contrast to the present work, mere cutting does not induce defence responses in *A. nodosum*, but actual grazing from the sea snail is required (Toth & Pavia 2000). Induction of defence mechanisms in response to water or air-borne cues has been reported for other brown macroalgae, too, as well as for red and green macroalgae, including the Ulvophyceaen alga *Ulva fenestrata* (Toth 2007; Yun et al. 2007; Thomas et al. 2011; Haavisto & Jormalainen 2019; Pereira et al 2020; van Alstyne et al. 2023). The nature of the cue is not always known. The previous studies have also often measured much slower responses (days) and longer distances (meters) than what was observed here, which may suggest that a different cue is involved.

Wounding releases various molecules to the surrounding media (for a review, see Aguilera et al. 2022). Often, production of different reactive oxygen species is also induced (e.g., Pohnert 2000; Ross et al. 2005). Some algae release toxins upon wounding (e.g., Paul & van Alstyne 1992), which, along with reactive oxygen species, can work as feeding deterrents (Collén et al. 1994; Küpper et al. 2001; McDowell et al. 2014; Hansel & Diaz 2021). In the Dasycladalean algae *Dasycladus vermicularis* and *A. acetabulum*, wounding has been shown to lead to accumulation of ATP, hydrogen peroxide and nitric oxide in the immediate vicinity of the wounding site (Torres et al. 2008). Furthermore, hydrogen peroxide production by *U. fenestrata* increases considerably when exposed to grazing from the sea snail *L. littorea* (Taenzer et al. 2024). However, in *D. vermicularis*, the production of reactive oxygen species increases 35–45 min after the wounding (Ross et al. 2005), which is too slow of a response to explain the present results, as the first signs of chloroplast aggregation were observed almost immediately after the wounding (Fig. S6). Interestingly, Torres et al. (2008) also showed that the addition of poorly hydrolysable analogues of ATP triggers a similar burst of reactive oxygen species as wounding and that by inhibiting algal purinoceptors, the burst is eliminated. Thus, in *A. acetabulum*, wounding leads to the release of ATP molecules to the media, which seems to induce the production of reactive oxygen species (Torres et al. 2008) and possibly also the aggregation of chloroplasts in nearby cells, observed here (Fig. 5), as a defence mechanism against attacks from sea slugs (Fig. 6).

In contrast to *A. acetabulum*, only local chloroplast movement responses were observed in *Bryopsis* sp. following cutting (Fig. 5). Bryopsidales algae, including *Caulerpa*, *Bryopsis* and *Halimeda* spp., are known to produce toxins such as caulerpenyne, kahalalide F and halimedatrial that are released into the media upon wounding, where they can be converted into even more toxic forms (Paul & van Alstyne 1992; Becerro et al. 2001; Jung & Pohnert 2001). These chemicals have been shown to repel herbivorous fish and have been proposed to contribute to the invasive nature of some of the Bryopsidales algae (e.g., Davis et al. 2005). Perhaps Bryopsidales algae rely on their chemical defences. However, it should be noted that the sea slug *Elysia rufescens*, which feeds on *Bryopsis* sp., tolerates kahalalide F and even uses it in its own defence (Becerro et al. 2001). In addition, Ulvophyceaen algae, such as *Ulva* spp., also possess chemical defences (e.g., van Alstyne et al. 2001).

Compared to *A. acetabulum*, *Bryopsis* sp. grows fast. Often, there is a trade-off between growth rate and chemical or physical defences against herbivory, observed e.g., in the unicellular green alga *Chlorella vulgaris* (Yoshida et al. 2004), the red macroalga *Laurencia dendroidea* (Sudatti et al. 2018) and the brown macroalga *Turbinaria ornata* (Bergman et al. 2016). Furthermore, Bryopsidales algae are coenocytic (multinucleate) and can regenerate from small, detached thallus parts (Kim et al. 2001). Predation by the sea slug *Lobiger serradifalci* was even shown to increase the dispersal of *C. taxifolia* (Žuljevic et al. 2001). On the contrary, an *A. acetabulum* cell has only a single nucleus, and herbivory can lead to the death of the individual (see Fig. S9). Bryopsidales algae might also respond to predation by increasing growth rates (Bulleri & Malquori 2015). To conclude, *Bryopsis* sp. may prioritise growth over protection and defence, while *A. acetabulum* needs to repair and prevent damages caused by both light and predators to survive.

## Acknowledgements

The work was supported by the Finnish Cultural Foundation (through the Foundation’s Post Doc Pool), Sakari Alhopuro Foundation and the Research Council of Finland (grant number 371558) to H.M., by the European Research Council under the European Union’s Horizon 2020 research and innovation program, grant agreement no. 949880 to S.C. (https://doi.org/10.3030/949880), and by Fundação para a Ciência e Tecnologia, grants 2020.03278.CEECIND to S.C. (https://doi.org/10.54499/2020.03278.CEECIND/CP1589/CT0012), CEECIND/01434/2018 to P.C. (https://doi.org/10.54499/CEECIND/01434/2018/CP1559/CT0003), CEECIND 04217/2020 to JWG, and UID/50017/2025 (https://doi.org/10.54499/UID/50017/2025) and LA/P/0094/2020 (https://doi.org/10.54499/LA/P/0094/2020) to CESAM.

