## Supplementary material for "Chloroplast movements in siphonous macroalgae in response to high light and grazing": Figs S1 to S9 and Tables S1 to S4

#### Supplementary figures

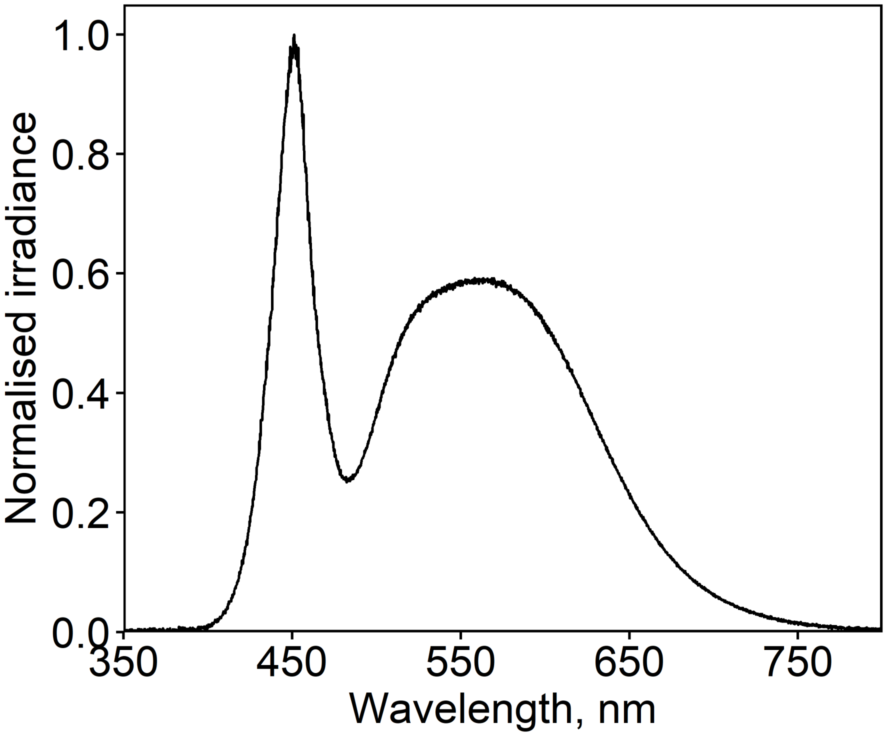
Normalised irradiance
1.0
0.8
0.6
0.4
0.2
0.0
350
450
550
650
750
Wavelength, nm

**Figure S1.** Normalized energy spectra of the white LED, used in the light treatments shown in Fig. 4.

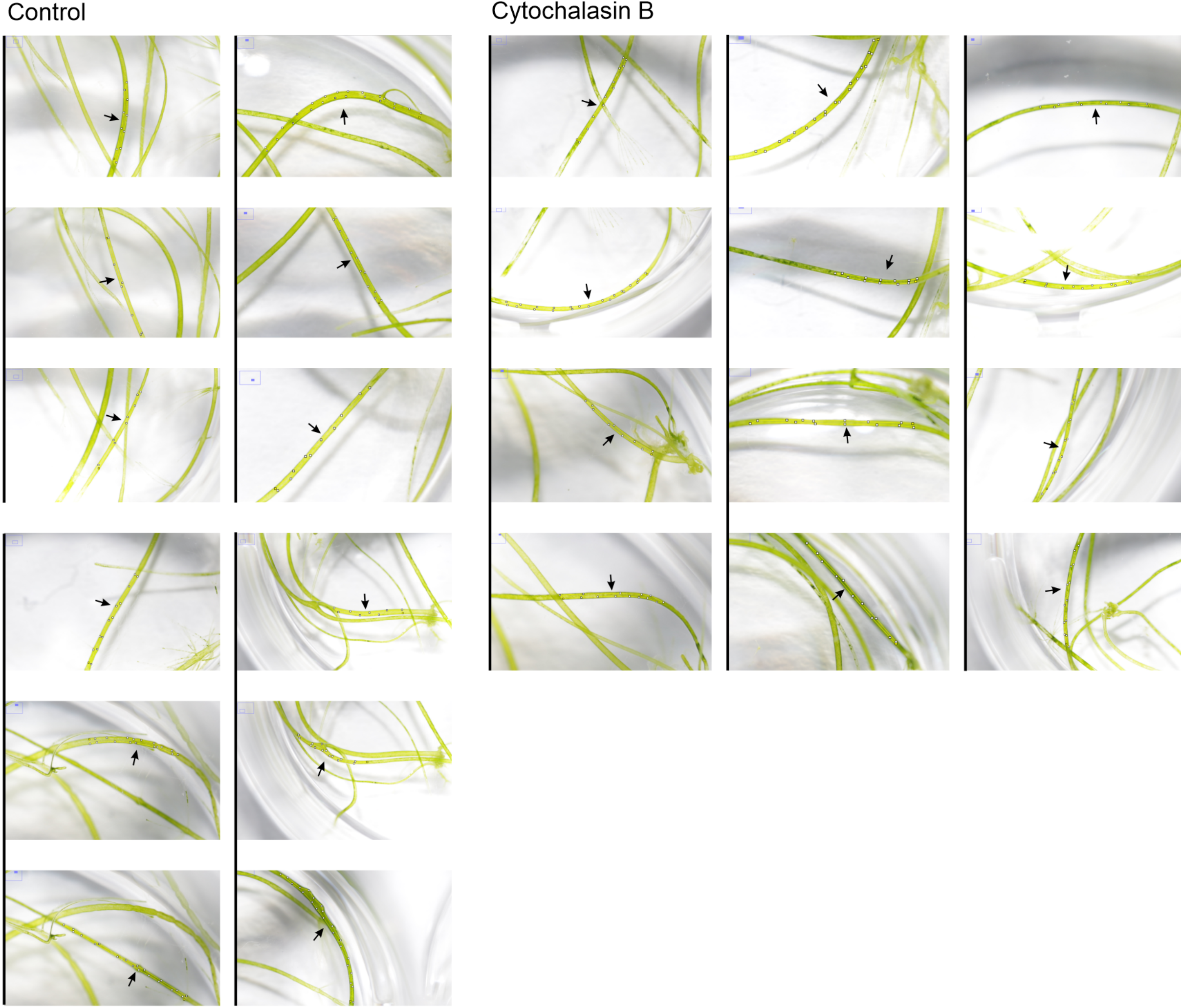
Control
Cytochalasin B

**Figure S2.** Selected areas of interest (AOI), each one from an individual *Acetabularia acetabulum* cell (12 biological replicates), selected from the captured RGB-images, used for pixel (the blue channel) analysis in Fiji for the high light experiment (Fig. 4), in the absence (Control) or presence of cytochalasin B, as indicated. The algal cell from which the AOI is selected, is highlighted with the black arrows. The AOIs are surrounded by thin yellow lines and small white boxes. The vertical lines highlight individual experiments (different photographs; four in the case of control and three in the case of cytochalasin B; note that the images above are cropped from the full photographs).

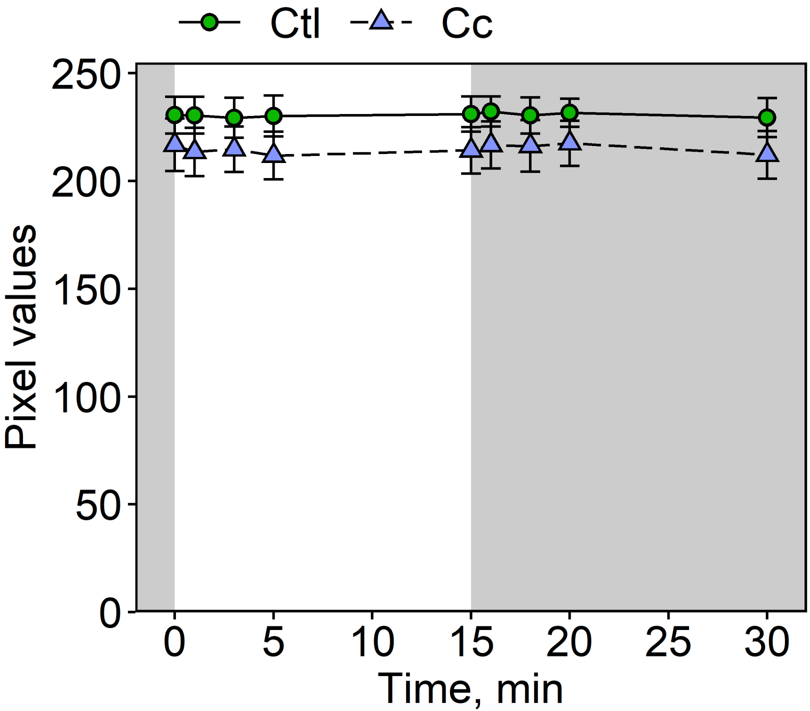

Ctl
Cc
Pixel values
0
50
100
150
200
250
0
5
10
15
20
25
30
Time, min

**Figure S3.** Quantification of the background signal from macro-images of *Acetabularia acetabulum* cells. Low-light-acclimated algae were placed in a well of a 4x6 well plate in 1 mL of artificial seawater and illuminated for 15 min with high light (illustrated with the white panel) and subsequently incubated in darkness for 15 min, in the absence (Ctl; circles) and presence of cytochalasin B (Cc; triangles). Areas containing no cells (only artificial seawater) were selected from the RGB-images and pixel values in the blue channel were quantified. Symbols show averages and error bars standard deviations based on 3 (Cc) or 4 (Ctl) technical replicates. See Fig. S3 for parts of the original images (showing both algal cells and the light background) and Fig. 4 for quantifications from algal parts.

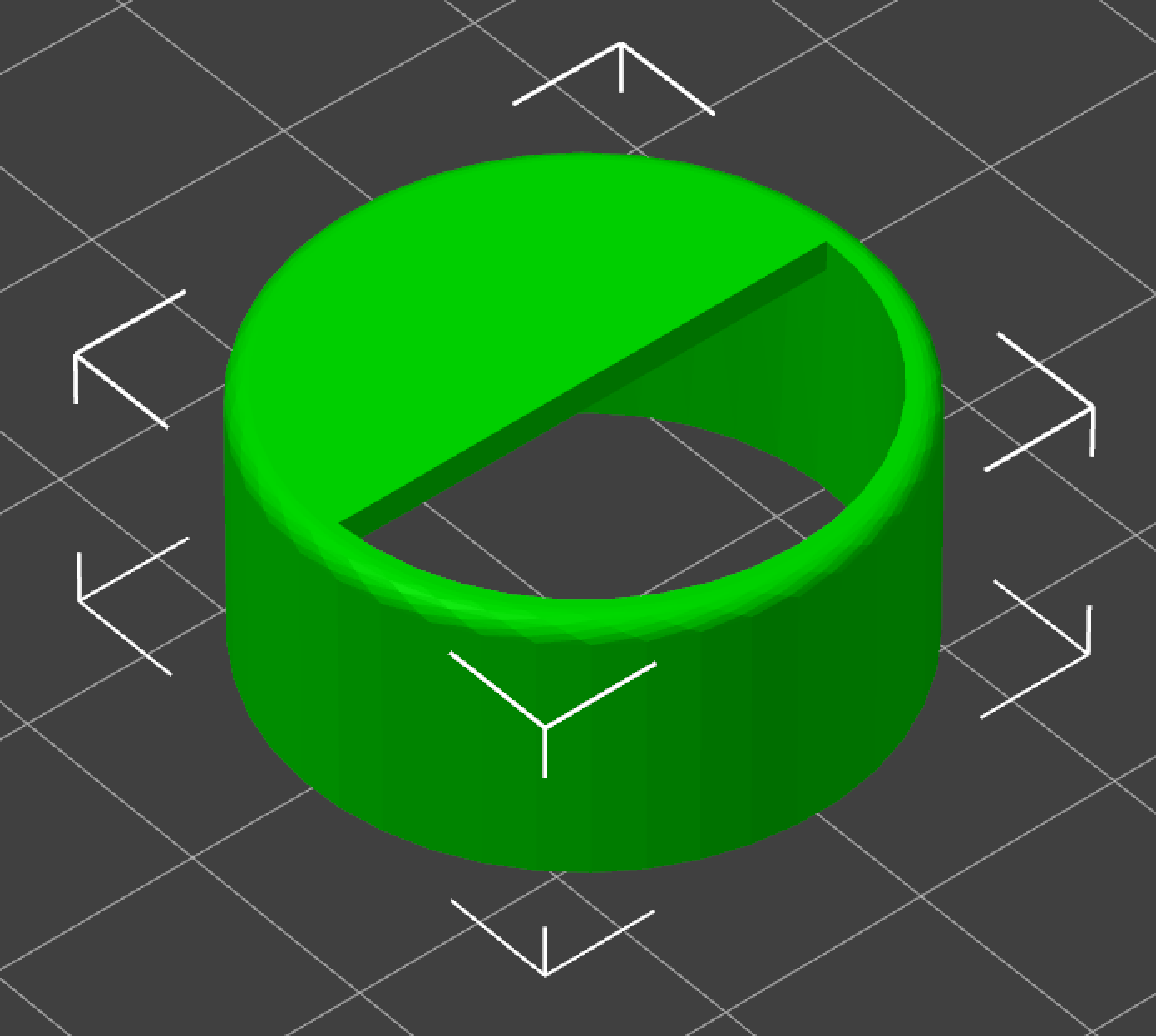

**Figure S4.** The model for a 3D-printed "half-shade" used to cover half of the well, in which the treated algae were placed, to create partial shade over the algal sample for the illumination treatments shown in Figs 2E–H and 3E–H.

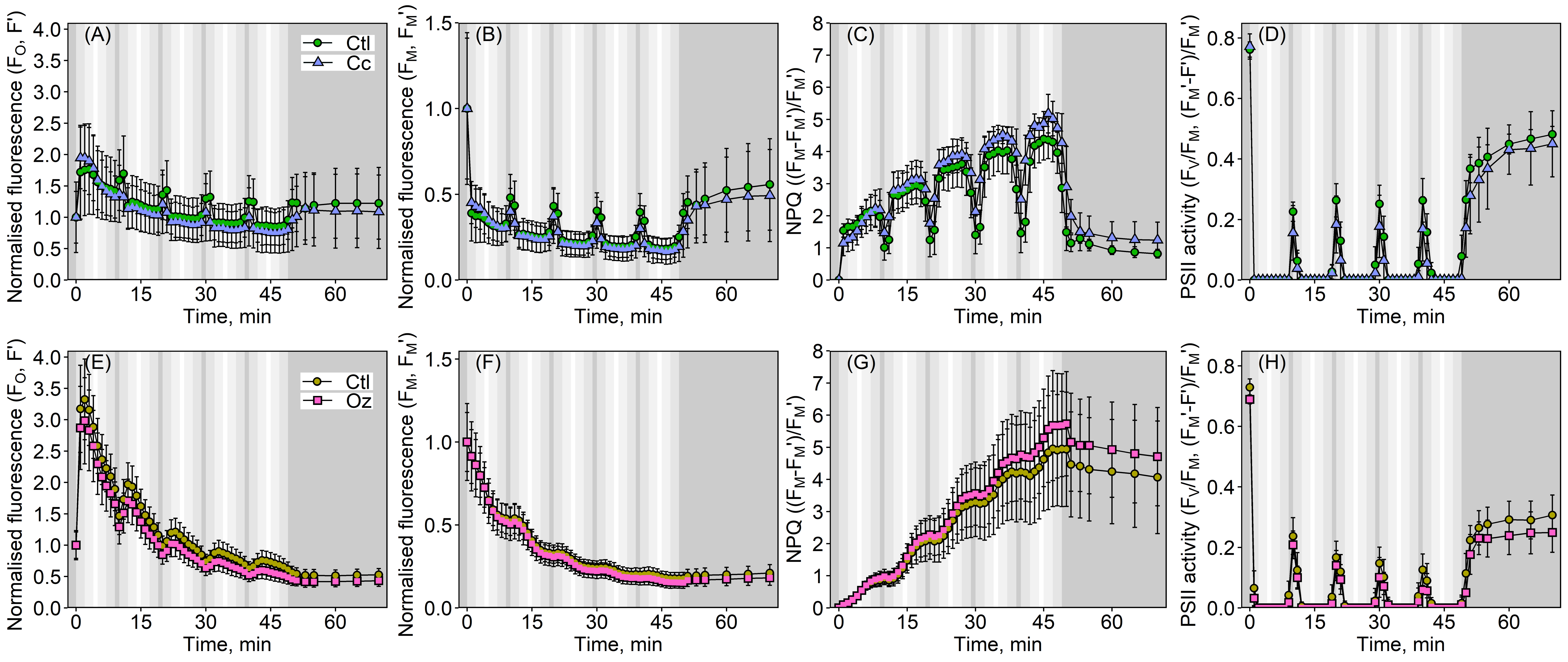

(A)
Ctl
Cc
Normalised fluorescence ($F_O$, F')
Time, min
(B)
Normalised fluorescence ($F_M$, $F_M$')
Time, min
(C)
NPQ (($F_M$-$F_M$')/$F_M$')
Time, min
(D)
PSII activity ($F_V$/$F_M$, ($F_M$'-F')/$F_M$')
Time, min
(E)
Ctl
Oz
Normalised fluorescence ($F_O$, F')
Time, min
(F)
Normalised fluorescence ($F_M$, $F_M$')
Time, min
(G)
NPQ (($F_M$-$F_M$')/$F_M$')
Time, min
(H)
PSII activity ($F_V$/$F_M$, ($F_M$'-F')/$F_M$')
Time, min

**Figure S5.** Effects of the inhibition of chloroplast movements on chlorophyll *a* fluorescence kinetics under gradually fluctuating light in *Acetabularia acetabulum* (A–D) *Bryopsis* sp. (E–H). Before the treatments, algae were dark acclimated for 20 min, in the absence (Ctl; circles) or presence of cytochalasin B (Cc; triangles) or oryzalin (Oz; squares). Algae were treated with periods of five min of increasing and five min of decreasing light (PPFD 0–1000 µmol $m^{-2}$ $s^{-1}$; visualized with the gradients of grey background colour), after which they were kept in darkness for 20 min. (A and E) Minimum fluorescence of dark-acclimated samples ($F_O$) or incident fluorescence under illumination (F'), normalised to the starting values. (B and F) Maximum fluorescence, obtained by firing saturating light pulses, of dark-acclimated samples ($F_M$) or under illumination ($F_M$'), normalised to the starting values. (C and G) Non-photochemical quenching (NPQ). (D and H) $F_V/F_M$ (dark-acclimated sample) or PSII operational yield (under illumination). The symbols show averages based on six to eight biological replicates and error bars show standard deviations.

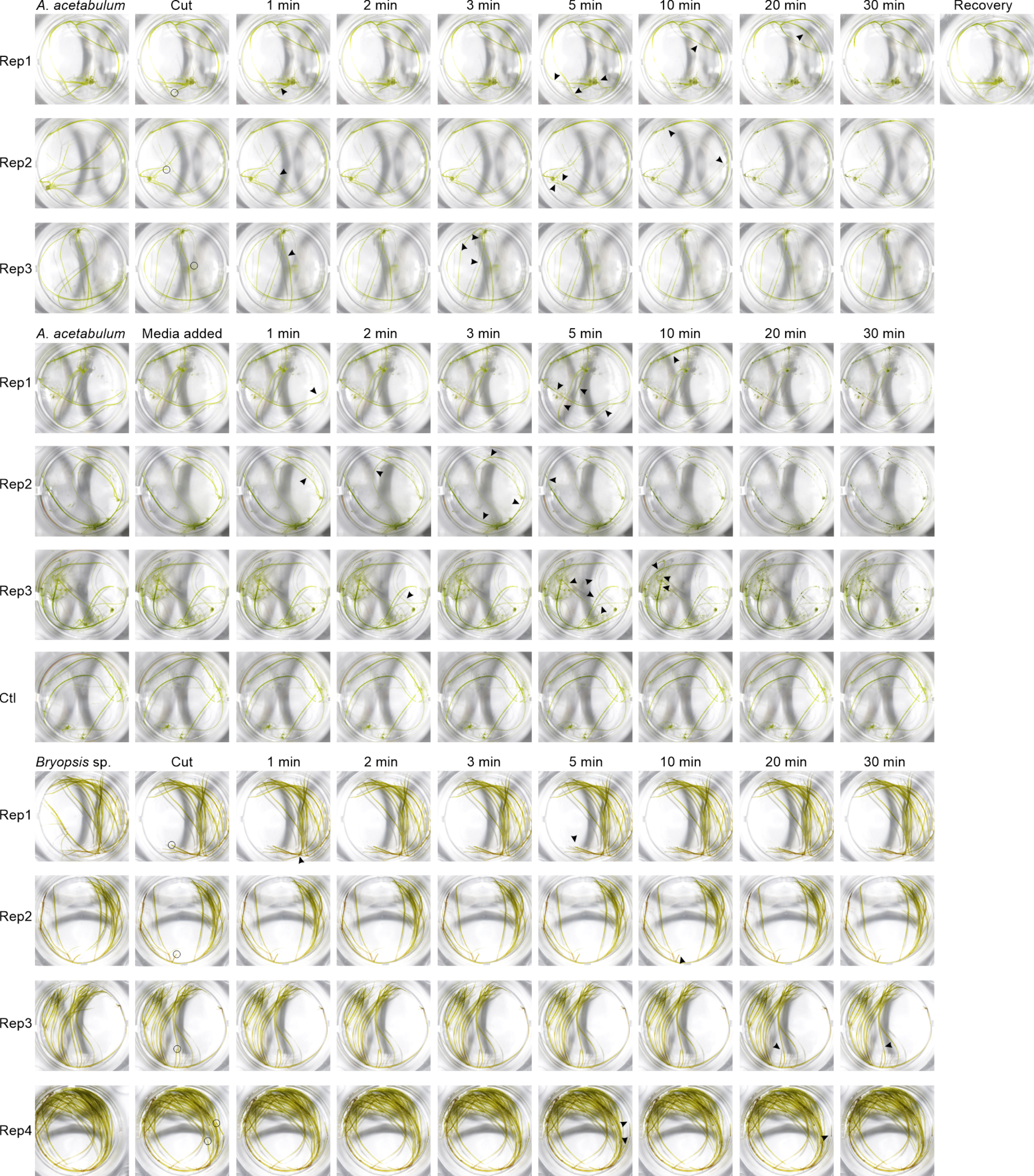
A. acetabulum
Cut
1 min
2 min
3 min
5 min
10 min
20 min
30 min
Recovery
Rep1
Rep2
Rep3
A. acetabulum
Media added
1 min
2 min
3 min
5 min
10 min
20 min
30 min
Rep1
Rep2
Rep3
Ctl
Bryopsis sp.
Cut
1 min
2 min
3 min
5 min
10 min
20 min
30 min
Rep1
Rep2
Rep3
Rep4

**Figure S6**. Macro-imaging of *Acetabularia acetabulum* and *Bryopsis* sp. cells, as indicated, in response to mechanical damage (cutting). Cells were placed in a well containing artificial seawater and imaged (first images; Rep1–4 indicate replications with different algae). A cell was then cut with scissors or, in the case of the second set of *A. acetabulum* images, 200 µl of artificial seawater medium was added, obtained from another well where *A. acetabulum* cells were cut to pieces right before the collection of the media (the ctl indicates a control (mock) experiment where only clean artificial seawater was added). Algae were imaged right after the cutting/addition of medium (second row of images; the black circles highlight the cutting sites) and after 1, 2, 3, 5, 10, 20 and 30 min, and after an overnight recovery (only the first replicate), as indicated. The arrowheads highlight first appearances of signs of chloroplast aggregation (i.e., darker or lighter spots).

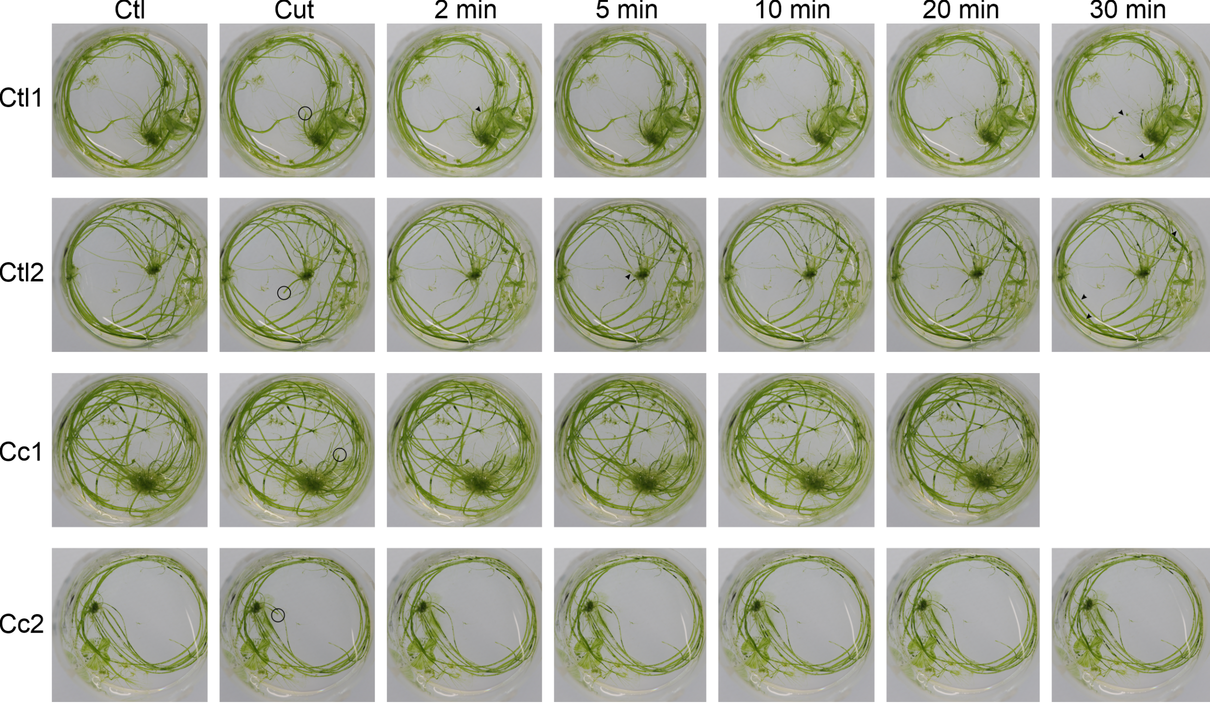
Ctl
Cut
2 min
5 min
10 min
20 min
30 min
Ctl1
Ctl2
Cc1
Cc2

**Figure S7**. Macro-imaging of *Acetabularia acetabulum*, in response to cutting, in the absence (Ctl) and presence (Cc) of cytochalasin B (added 20 min prior to the imaging). Low-light-acclimated cells were placed in a well containing artificial seawater and imaged (first images; numbers 1 and 2 indicate replications with different algae). A cell was then cut with scissors (the cutting site is indicated with circles; the second images) and algae were imaged after 2, 5, 10, 20 and 30 min, as indicated. The black arrowheads highlight first appearances of signs of new chloroplast aggregation (i.e., darker or lighter spots).

(A)

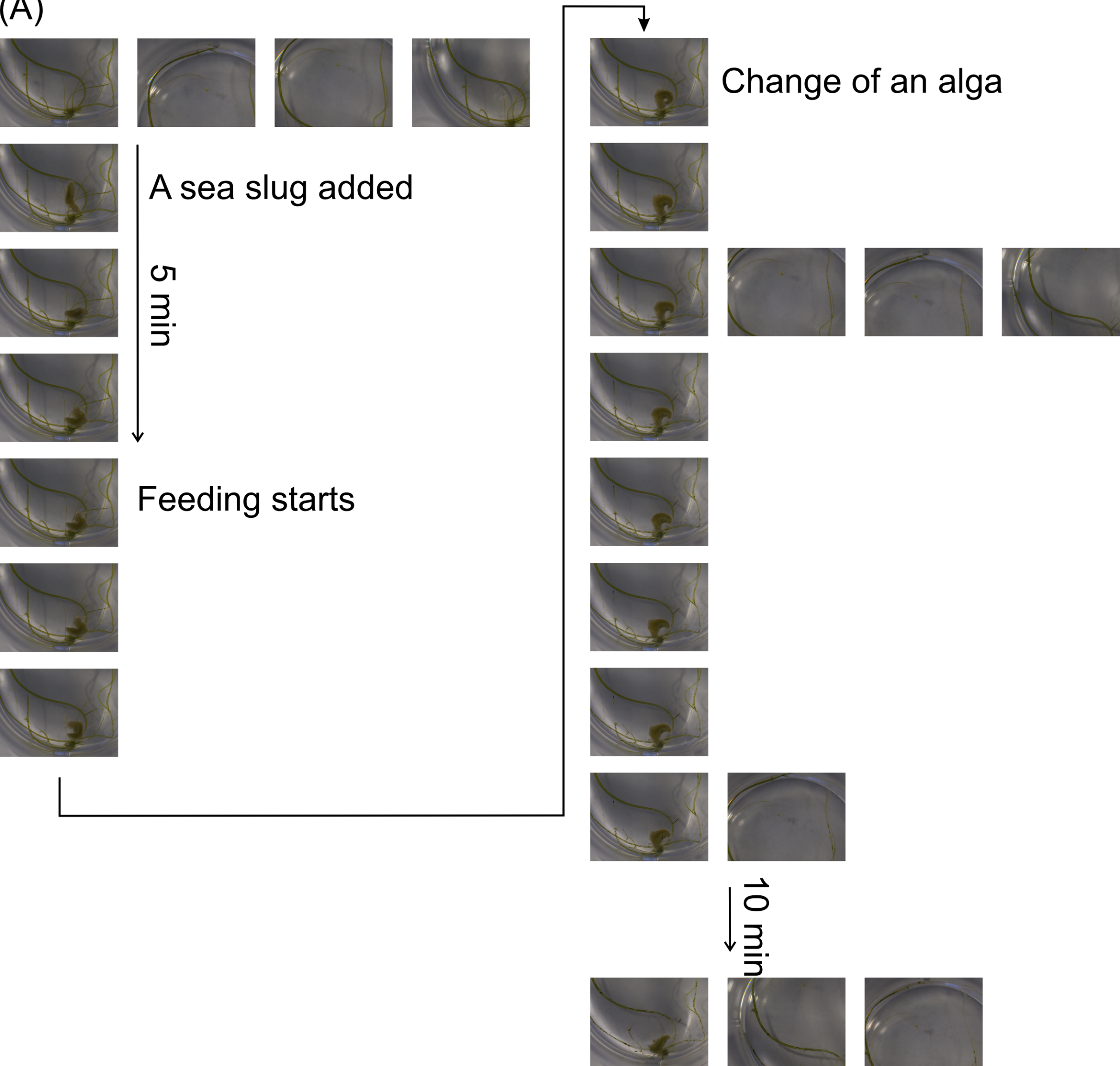

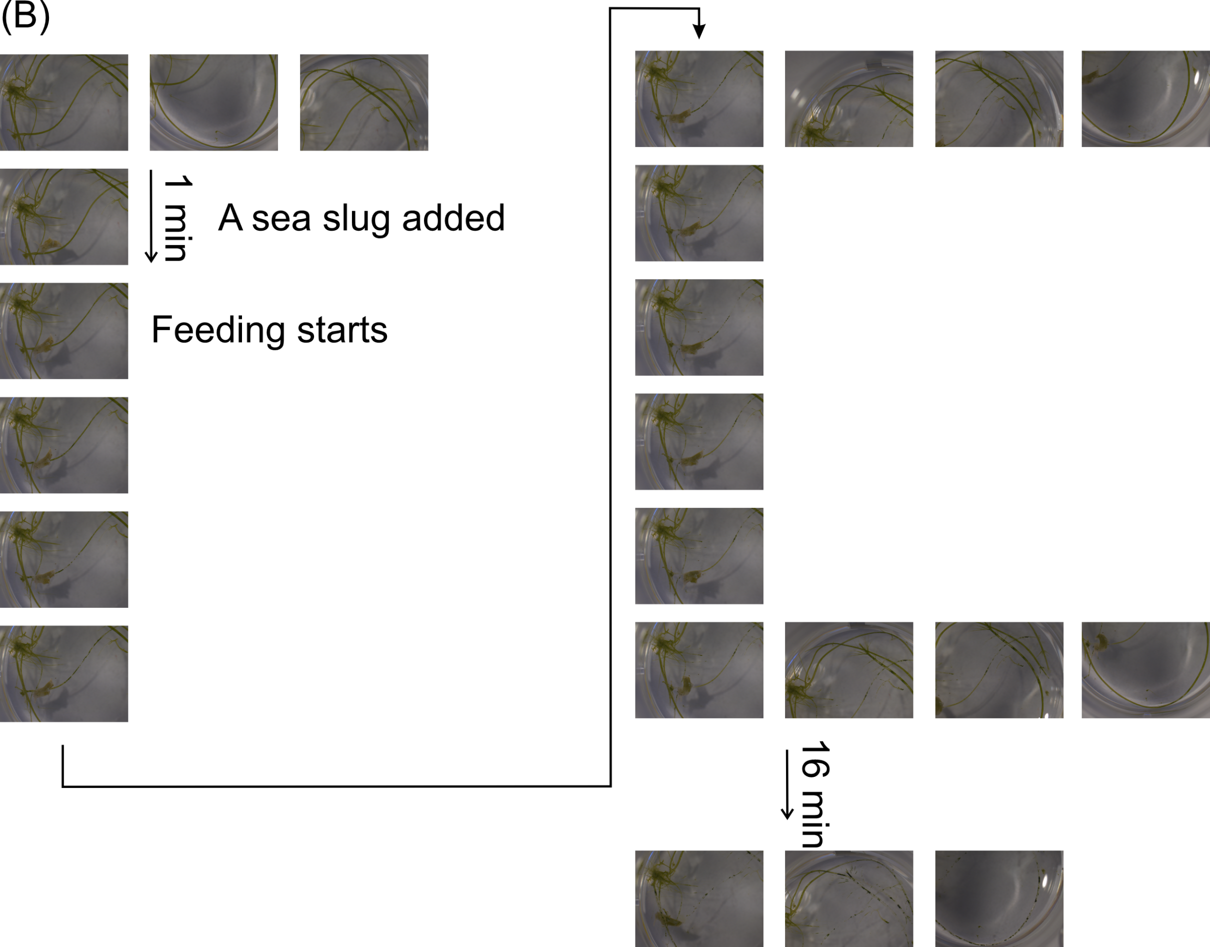

(B)
1 min
A sea slug added
Feeding starts
16 min

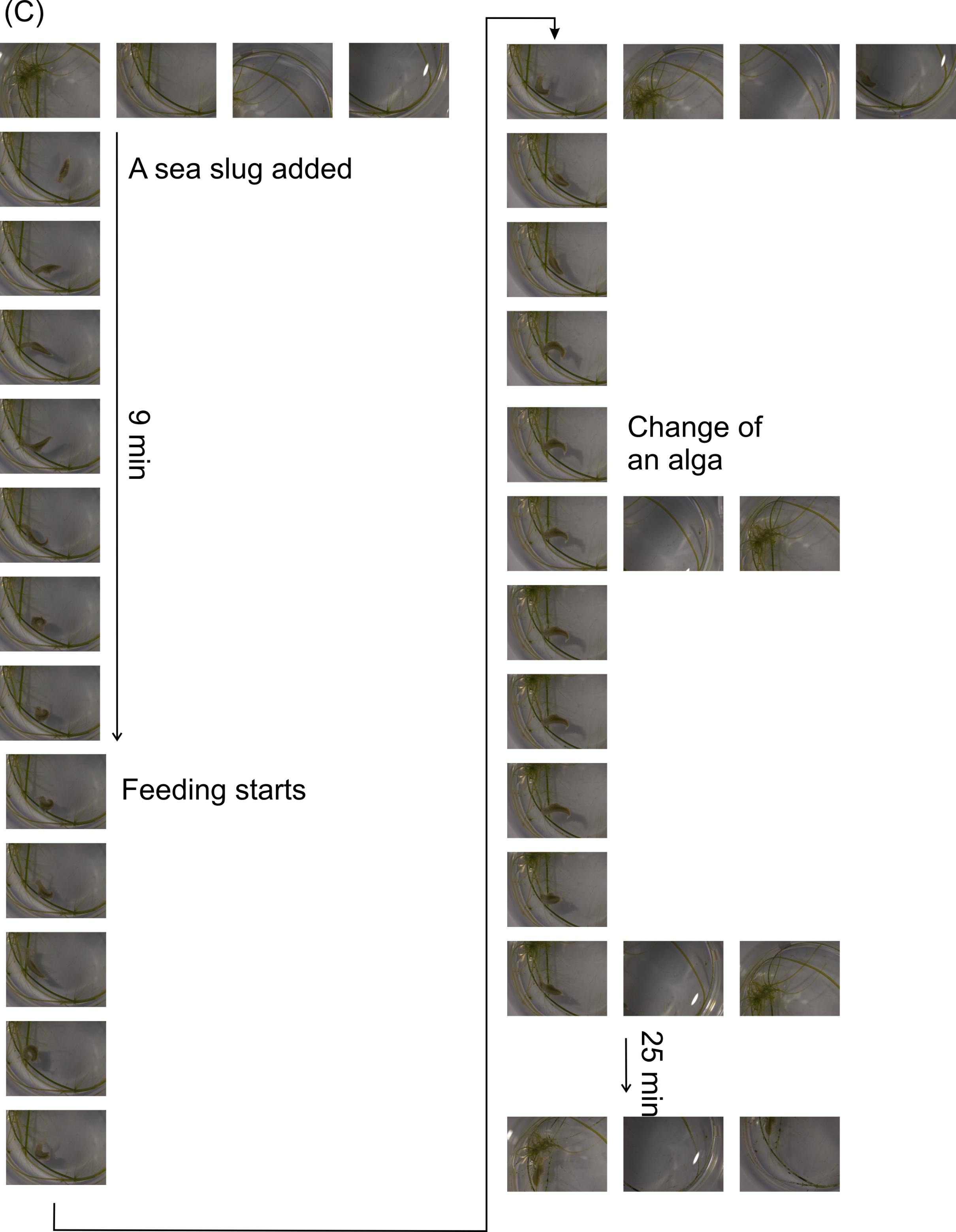
(C)
A sea slug added
9 min
Feeding starts
Change of
an alga
25 min

**Fig. S8**. An *Elysia timida* sea slug feeding on *Acetabularia acetabulum*. A sea slug (highlighted with the circle) was placed in a well with intact algae. The feeding started (according to visual inspection) after the indicated times. Imaging was then repeated every 1 min, unless otherwise indicated. The 3 vertical rows of images represent three individual replications with different algae and sea slugs. The first four to three vertical images show the intact algae prior the addition of the slug. (A–C) Biological replicates.

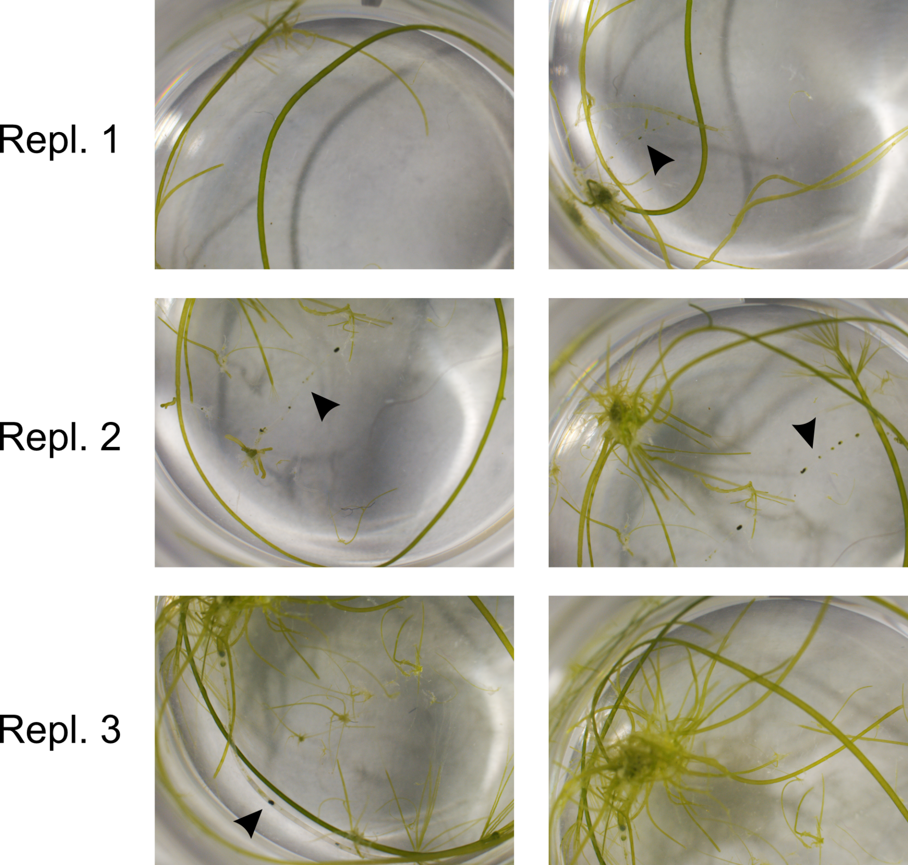
Repl. 1
Repl. 2
Repl. 3

**Fig. S9**. *Acetabularia acetabulum* cells, imaged after 3 days of recovery under lowlight, after feeding by *Elysia timida* sea slugs (Fig. S8). Three individual replications (Repl. 1–3) are shown, each imaged twice. The black arrows indicate the alga the sea slug was feeding on (before it was removed and the algae let to recover). Brightness and contrast have been adjusted equally in all images. To see the feeding data, see Figs 6 and S8.

### Supplementary tables

**Table S1**. Results of the Type III ANOVA from the linear mixed model on the effect of the treatment (control without cytochalasin B, ctl; the presence of cytochalasin B; cc), time (a dark-light-dark-series) and their interaction on chloroplast density (probed by calculating mean pixel intensity from RGB-images) in *Acetabularia acetabulum*. See Fig. 4A for the data. NumDF = Numerator Degrees of Freedom; DenDF = Denominator Degrees of Freedom (Satterwaite estimated degrees of freedom).

| **Effect (mean)** | **Sum Sq** | **Mean Sq** | **NumDF** | **DenDF** | **F value** | **Pr(>F)** |
|---|---|---|---|---|---|---|
| **Treatment (ctl vs cc)** | 0.10356 | 0.103562 | 1 | 22 | 15.7062 | 0.0006599 |
| **Time** | 0.20705 | 0.025881 | 8 | 176 | 3.9251 | 0.0002722 |
| **Treatment:Time** | 0.28192 | 0.035240 | 8 | 176 | 5.3445 | 5.102e-06 |

**Table S2**. Results of the Type III ANOVA from the linear mixed model on the effect of the treatment (control without cytochalasin B, ctl; the presence of cytochalasin B, cc), time (a dark-light-dark-series) and their interaction on chloroplast distribution((probed by calculating standard deviation in pixel intensity from RGB-images) in *Acetabularia acetabulum*. See Fig. 4B for the data. NumDF = Numerator Degrees of Freedom; DenDF = Denominator Degrees of Freedom (Satterwaite estimated degrees of freedom).

| **Effect (standard deviation)** | **Sum Sq** | **Mean Sq** | **NumDF** | **DenDF** | **F value** | **Pr(>F)** |
|---|---|---|---|---|---|---|
| **Treatment (ctl vs cc)** | 0.17418 | 0.174181 | 1 | 22 | 5.6149 | 0.0270012 |
| **Time** | 0.66615 | 0.083269 | 8 | 176 | 2.6843 | 0.0082825 |
| **Treatment:Time** | 1.04181 | 0.130226 | 8 | 176 | 4.1980 | 0.0001268 |

**Table S3**. Results of the Type III ANOVA from the linear mixed model on the effect of the treatment (cutting; addition of medium collected from the vicinity of cut algae, media; addition of clean artificial seawater, mock), time and their interaction on chloroplast density (probed by calculating mean pixel intensity from RGB-images) in *Acetabularia acetabulum*. See Fig. 5A for the data. NumDF = Numerator Degrees of Freedom; DenDF = Denominator Degrees of Freedom (Satterwaite estimated degrees of freedom).

| **Effect (mean)** | **Sum Sq** | **Mean Sq** | **NumDF** | **DenDF** | **F value** | **Pr(>F)** |
|---|---|---|---|---|---|---|
| **Treatment (cutting, media, mock)** | 2.9246 | 1.46231 | 2 | 13 | 8.6348 | 0.0041123 |
| **Time** | 7.9147 | 0.98933 | 8 | 104 | 5.8419 | 3.693e-06 |
| **Treatment:Time** | 7.7495 | 0.48434 | 16 | 104 | 2.8600 | 0.0006889 |

**Table S4**. Results of the Type III ANOVA from the linear mixed model on the effect of the treatment (addition of medium collected from the vicinity of cut algae, media; or addition of clean artificial seawater, mock), time and their interaction on chloroplast distribution (probed by calculating standard deviation in pixel intensity from RGB-images) in *Acetabularia acetabulum*. See Fig. 5B for the data. NumDF = Numerator Degrees of Freedom; DenDF = Denominator Degrees of Freedom (Satterwaite estimated degrees of freedom).

| Effect (standard deviation) | Sum Sq | Mean Sq | NumDF | DenDF | F value | Pr(>F) |
|---|---|---|---|---|---|---|
| **Treatment (cutting, media, mock)** | 1.3949 | 0.69743 | 2 | 13 | 5.7807 | 0.01599 |
| **Time** | 19.1848 | 2.39810 | 8 | 104 | 19.8769 | < 2.2e-16 |
| **Treatment:Time** | 8.4957 | 0.53098 | 16 | 104 | 4.4011 | 1.546e-06 |
